# Stimulation of the *Caulobacter crescentus* surface sensing pathway by deletion of a specialized minor pilin-like gene

**DOI:** 10.1101/2025.02.12.637803

**Authors:** Farah Obeid Charrouf, Gregory B Whitfield, Courtney K Ellison, Yves V Brun

**Affiliations:** Département de Microbiologie, Infectiologie et Immunologie, Université de Montréal, 2900 Boulevard Édouard-Montpetit, Montréal, Québec H3T 1J4, Canada; Department of Microbiology, University of Georgia, Athens, GA, USA

## Abstract

Bacteria colonize surfaces through complex mechanisms of surface sensing. Pili are dynamic bacterial appendages that play an important role in this process. In *Caulobacter crescentus*, tension on retracting, surface-bound pili triggers the rapid synthesis of the adhesive holdfast, which permanently attaches cells to surfaces. However, the detailed mechanisms of pilus-mediated surface sensing are unclear. In this study, we used a genetic screen to isolate mutants with altered pilus activity to identify genes that may be involved in pilus-mediated surface-sensing. This screen identified *cpaL*, whose deletion led to reduced piliation levels, and surprisingly, a threefold increase in surface adhesion due to increased holdfast production. To understand this finding, we compared holdfast synthesis in wild-type and *cpaL* mutant cells under conditions that block pilus retraction. While this treatment increased holdfast production in wild-type cells by triggering the surface-sensing pathway, no increase was observed in the *cpaL* mutant, suggesting that mutation of *cpaL* maximally stimulates surface-sensing. Furthermore, when the *cpaL* mutant was grown in a medium that blocks the surface sensing pathway, cells exhibited decreased surface attachment and holdfast production, consistent with a role for CpaL in pilus-dependent surface sensing in *C. crescentus*. To better understand the function of CpaL, we analyzed its predicted structure, which suggested that CpaL is a minor pilin fused to a mechanosensitive von Willebrand factor type A (vWA) domain that could be accommodated at the pilus tip. These results collectively position CpaL as a strong candidate for a mechanosensory element in pilus-mediated surface sensing.

**Importance:** Surface sensing is a crucial mechanism that allows bacteria to change their behaviors to adapt to life on a surface. Surface recognition by bacteria is the initial step toward surface colonization and biofilm formation. In *Caulobacter crescentus*, tight adherence (Tad) pili play a key role in surface recognition and adaptation. However, the mechanism of pilus-mediated surface sensing and the proteins that influence this process remain unknown. Here, we demonstrate that CpaL, a potential pilus tip mechanosensory protein, could be the major element of Tad pilus-mediated surface attachment and colonization in *C. crescentus*. CpaL plays an important role in the regulation of holdfast synthesis and production upon surface contact. By identifying CpaL as a key player in the process of surface recognition, our work offers valuable insights into the mechanisms of bacterial adhesion.

## Introduction

Surface recognition and attachment are essential steps in the process of bacterial surface colonization to form biofilms. Biofilms are cohesive, multicellular microbial communities wherein resident bacteria are protected from harsh environmental conditions, including variations in osmolarity, pH, nutrient accessibility, shear forces, or exposure to antibacterial agents (1, 2). Bacteria are known to sense mechanical or physical cues upon contact with solid substrata, allowing them to respond with phenotypic changes that transform free-swimming planktonic bacterial cells into surface-adherent cells that form biofilms (3–5). The process by which bacterial cells sense a surface via mechanical stimuli and convert this signal into downstream cellular processes is called surface-sensing (6, 7). A great deal of effort has been expended in deciphering the mechanisms of surface sensing in bacteria, establishing a pivotal role for extracellular appendages such as flagella and pili in this process (8–11).

Pili are thread-like, proteinaceous appendages that physically interact with their surroundings and are critical for many important cellular processes such as adherence, aggregation, biofilm formation, horizontal gene transfer, and virulence in some pathogenic species (12, 13). Pili are composed of thousands of repeats of a small protein subunit, the major pilin, and less abundant minor pilin(s) (12, 14). A subset of pili, called type IV pili (T4P), are characterized by their dynamic activity, exhibiting the ability to extend away from the cell surface and subsequently retract back into the bacterial cell by polymerization and depolymerization of the pilus fiber, respectively. This is achieved by a complex membrane-spanning machine that draws from a pool of pilins in the inner membrane to assemble the fiber, which passes through an outer membrane channel (14). Based primarily on differences in the motor components of the T4P machinery, recent phylogenetic analyses have divided the T4P into three subclasses: T4aP, T4bP, and T4cP (11, 15, 16). Among these, T4cP, also known as tight adherence (Tad) pili, are thought to have evolved from an archaeal ancestor. Tad pili are broadly distributed among bacteria, including in the freshwater bacterium *Caulobacter crescentus*, where they have been implicated in adhesion and surface-sensing (11, 17).

*C. crescentus* exhibits a dimorphic life cycle, wherein each cell division produces a nonmotile stalked cell and a motile swarmer cell harboring multiple Tad pili and one flagellum at the same pole (18). The swarmer cell subsequently undergoes differentiation into a stalked cell as it developmentally progresses through the cell cycle, whereupon it synthesizes a compositionally complex adhesin called holdfast, which mediates permanent surface attachment (10, 18, 19). Interestingly, surface contact by a swarmer cell can hasten the differentiation process by rapidly triggering multiple processes, starting with the retraction of the pili into the cell, cessation of further pilus activity, ejection of the flagellum, and finally synthesis of the holdfast (11). The rapid, surface-contact mediated production of holdfast requires the presence of the pilus, and physical obstruction of pilus retraction leads to rapid holdfast synthesis even in the absence of surface contact, implying that tension on retracting, surface-bound pili may be a cue to sense surface contact (11, 20). However, the mechanistic details of how Tad pili sense surface contact, convey the signal across the cell envelope, and translate it into an output capable of upregulating surface-associated behaviors, remain poorly understood.

In this study, we address these knowledge gaps in our mechanistic understanding of Tad pilus-mediated mechanosensing. We performed a genetic screen that identified mutations in *cpaL*, whose deletion led to significantly reduced pilus synthesis. However, despite reduced pilus production, we found that this mutant showed increased surface adherence and holdfast synthesis. Stimulation of holdfast production was not observed when a *cpaL* mutant was cultured in a defined medium that nutritionally restricts surface sensing. In addition, holdfast synthesis did not increase when pilus retraction was blocked in the *cpaL* mutant to constitutively stimulate surface sensing. Finally, we examined the predicted structure of CpaL using AlphaFold3 and found that CpaL is comprised of a pilin-like module connected to a von Willebrand factor type A (vWA) domain, a fold that is often implicated in mechanosensing. Further modelling suggests that CpaL could form a complex with the minor pilins CpaJ and CpaK, that can be accommodated at the distal end of the pilus. Collectively, these data suggest that CpaL plays an important role in the pilus-dependent surface sensing pathway of *C. crescentus*.

## Results

### A screen for mutations conferring resistance to the pilus-dependant phage ΦCbK identifies the *cpaL* gene

Tad pili exhibit dynamic cycles of extension and retraction, which play a crucial role in surface-sensing in *C. crescentus* (11). To understand this process, we sought to identify genes in *C. crescentus* that impact pilus activity using a forward genetics approach to identify mutants that are resistant to the pilus-dependent phage ΦCbK. We reasoned that impaired pilus activity, e.g. through impaired biosynthesis or dynamics, might increase resistance to the phage ΦCbK. To carry out our screen and all subsequent analyses, we used *C. crescentus pilA-cys* as the parent background, where WT cells are modified to incorporate a cysteine residue in the major pilin PilA, allowing for pilus labeling and modulation (11, 21). A pooled transposon mutant library was generated in the parent strain and then mixed with ΦCbK, and phage resistant mutants were isolated. The ability of these mutants to elaborate a pilus was analyzed via microscopy by labeling pili with a thiol-reactive, maleimide-conjugated fluorophore. Mutants that produced pili and yet were ΦCbK-resistant were chosen for further study, and the site of transposon insertion was determined (Figure S1) (11, 21). Among these mutants, five transposon insertions mapped to *cpaL* (*CCNA_0199*) (Figure S1). The gene *cpaL* is located outside the main pilus gene cluster and has previously been shown to contribute to phage sensitivity in *C. crescentus* (22).

We generated an unmarked, in-frame deletion of *cpaL* and assessed the sensitivity of this mutant to ΦCbK. We spotted serial dilutions of ΦCbK onto growth plates with *C. crescentus* incorporated into the top agar and analyzed the formation of plaques due to phage infection (Figure 1A). The phage-susceptible parent strain exhibited the formation of clear plaques up to the 10^-5^ phage dilution, while the pilus-deficient mutant Δ*pilA* displayed complete resistance to ΦCbK. In contrast, the Δ*cpaL* mutant demonstrated intermediate resistance to ΦCbK, with formation of cloudy plaques visible only up to the 10^-1^ phage dilution. The ΦCbK resistance phenotype in Δ*cpaL* was completely reversed when complemented with a wild-type copy of *cpaL* on a replicating plasmid (Figure 1A). The growth curves of all the analyzed strains were comparable, indicating that ΦCbK resistance in the Δ*cpaL* mutant is not due to a variation in growth rate (Figure S2). Together, these results indicate that *cpaL* plays an important role in pilus biosynthesis.

**Figure 1:**
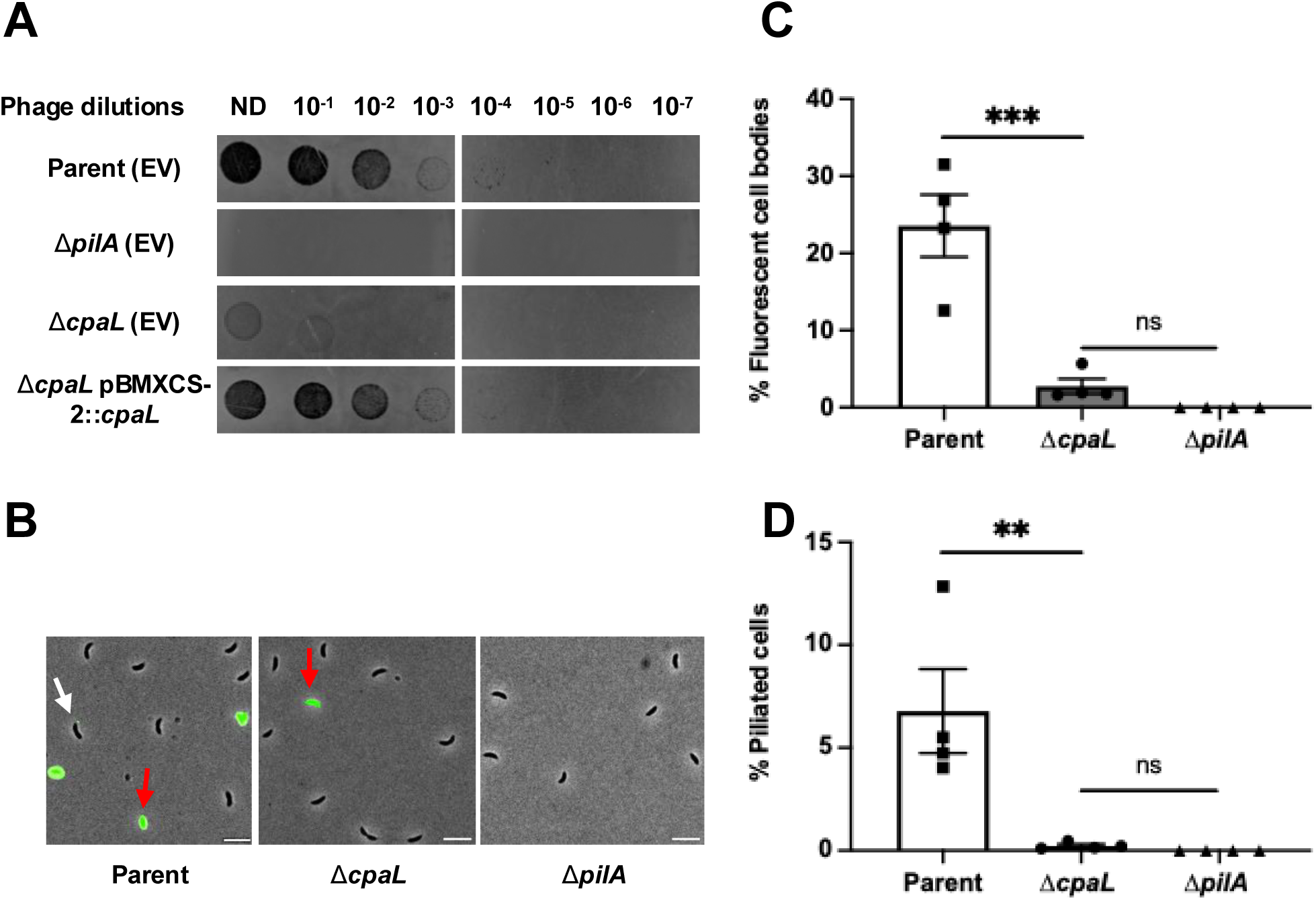
Deletion of *cpaL* reduces sensitivity to the pilus-specific phage ΦCbK by reducing pilus activity. **A:** Top agar phage sensitivity assays for the parent strain (NA1000 *pilA*-cys), Δ*pilA* (phage-resistant strain lacking pili), the Δ*cpaL* mutant, and the plasmid-complemented Δ*cpaL* mutant. Serial dilutions of the phage were spotted on plates with top agar containing each bacterial strain. ND, no dilution; EV, empty vector. **B:** Representative microscopy images for synchronized swarmer cells of the parent strain (NA1000 *pilA*-cys) and the isogenic Δ*cpaL* mutant, labeled with the AF488-maleimide dye (green) that reacts with the engineered cysteine residue in the major pilin, PilA. Scale bars, 10 μm. White arrows indicate cells with labeled pili, red arrows indicate fluorescent cell bodies. **C:** Quantification of the percentage of cells with fluorescent cell bodies in synchronized populations of the indicated strains labelled with AF488-maleimide. **D:** Quantification of the percentage of piliated cells in synchronized swarmer cells of the indicated strains labelled with AF488-maleimide dye (green). Results are the mean of four independent biological replicates, with at least 500 cells analyzed per replicate. Error bars represent the standard error of the mean (SEM). Statistical comparisons were made using Tukey’s multiple comparisons test. ***, P <0.001; **, P < 0.01; ns, no significant difference.

### The Δ*cpaL* mutant produces fewer pili per cell, but with an increased average length

Since the Δ*cpaL* mutant was less sensitive to the pilus-dependent phage ΦCbK, we hypothesized that this could be due to a decrease in the amount of pilus activity within the population. To visualize pilus activity, we observed the internalization of externally labeled pilins into the cell during pilus retraction, which results in cells with fluorescent bodies in *C. crescentus* (Figure 1B) (11). We quantified the proportion of fluorescent cell bodies among synchronized swarmer cells stained for pili, comparing the parent and mutant strains. Approximately 23.5% of synchronized swarmer cells of the parent strain had fluorescent cell bodies after labeling, consistent with previous reports (11, 23). In contrast, less than 3% of the Δ*cpaL* synchronized swarmer cells displayed fluorescent cell bodies (Figure 1B and C). Moreover, approximately 6% of parent cells had visible pili, whereas piliated cells were rarely observed for the Δ*cpaL* mutant (Figure 1B and D). The *ΔpilA* mutant lacking pili served as a negative control, showing no fluorescence. These results indicate that the Δ*cpaL* mutant exhibits significantly less pilus activity than the parental strain, but that those pili that are produced can retract. The dynamic activity of pili produced by the Δ*cpaL* mutant was examined by time-lapse microscopy, which revealed a distribution of dynamic behaviors that is consistent with what has been observed previously for *C. crescentus* (Movies S1-S10) (11).

Next, we incubated synchronized parental and Δ*cpaL* swarmer cells with PEG5000-maleimide (PEG5000-mal) to block pilus retraction, through its reaction with the modified PilA cysteine residue, while simultaneously labeling pili before imaging, as described previously (11). We found that around 53% of the parental swarmer cell population exhibited piliated cells (Figure 2A and B). In contrast, around 6% of swarmer cells from the Δ*cpaL* mutant population produced pili (Figure 2A and B). Finally, the production of pili was restored to parental levels when the Δ*cpaL* mutant was complemented with plasmid-borne wild-type *cpaL* (Figure S3).

**Figure 2:**
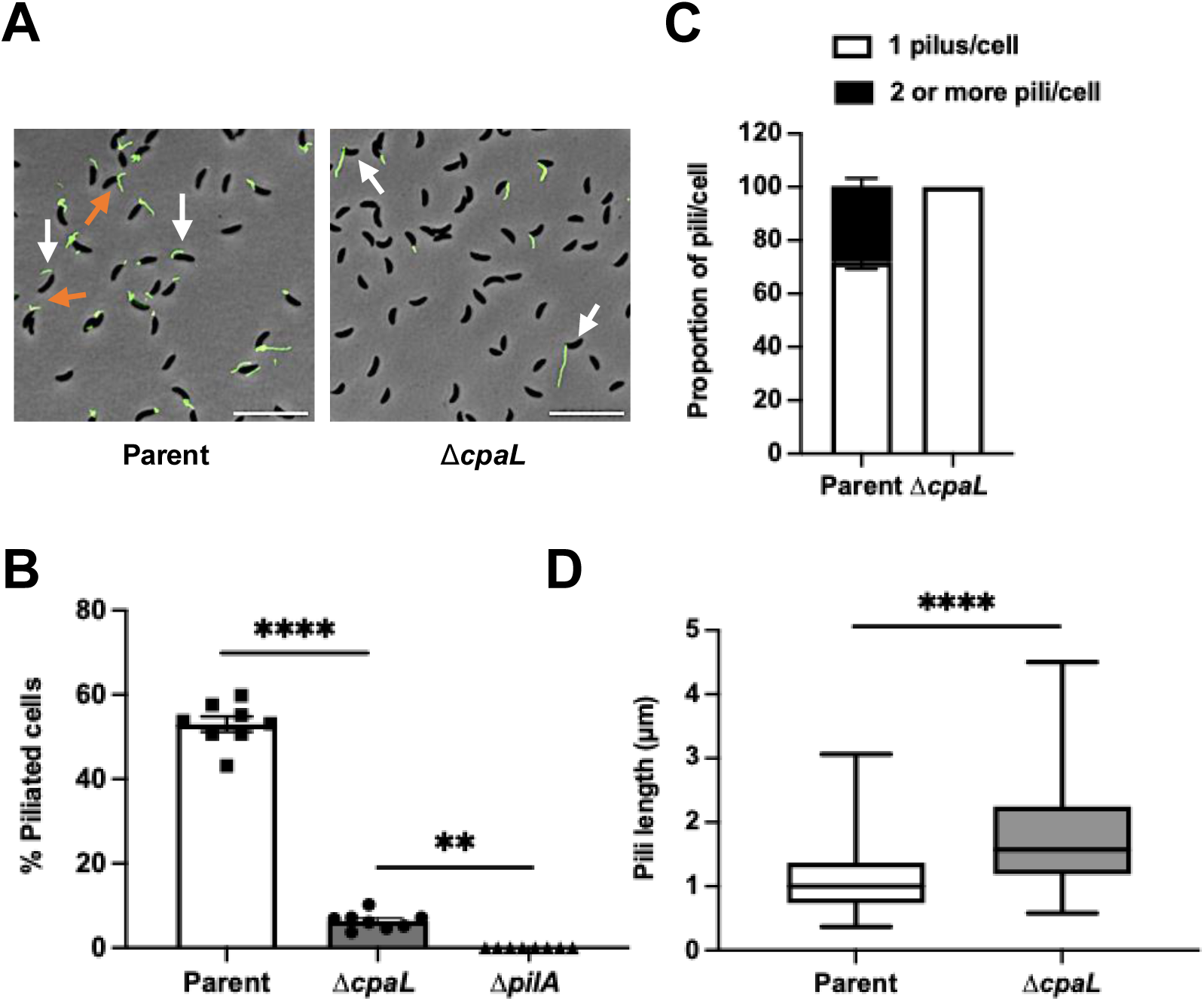
Deletion of *cpaL* reduces the number of pili produced per cell while increasing their average length. **A:** Representative microscopy images of synchronized swarmer cells of the parent strain (NA1000 *pilA*-cys) and the isogenic Δ*cpaL* mutant labeled with AF488-maleimide (green) that reacts with the engineered cysteine residue in the major pilin, PilA, in conjunction with pilus retraction blocking using PEG5000-mal, which reacts with the same cysteine residue. White arrows indicate cells with a single pilus per cell. Orange arrows indicate cells with two or more pili per cell. Scale bars, 10 μm. **B:** Quantification of the percentage of piliated cells in synchronized populations of the indicated strains when pilus retraction was blocked with PEG5000-mal. Results are the mean of eight independent biological replicates, with at least 200 cells analyzed per replicate. Error bars represent the standard error of the mean. Statistical comparisons were made using Tukey’s multiple comparisons test. **C:** Quantification of the number of pili produced per piliated cell of the indicated strains, measured in synchronized populations after blocking pilus retraction with PEG5000-mal. Results are the mean of four independent biological replicates with at least 200 cells analyzed per replicate. Error bars represent the standard error of the mean. **D:** Average length of pili produced by synchronized swarmer cells of the indicated strains after blocking with PEG5000-mal. 100 pili were measured for each strain. Error bars indicate the minimum to maximum range of lengths. Statistical comparison was made using a two-tailed unpaired T-test. ****, P <0.0001; **, P < 0.01.

Next, we analyzed the number of pili produced by each piliated cell after blocking and found that the Δ*cpaL* mutant produced only one pilus per piliated cell, in contrast to the parent strain, where 29% of piliated cells elaborated more than one pilus (Figure 2A and C). In addition, the single pilus produced by the Δ*cpaL* mutant had an average length of 1.81 µm, which is significantly longer than the pili produced by the parent strain (1.02 µm; consistent with previous reports) (11) (Figure 2A and D). These findings reinforce the importance of CpaL for Tad pilus biosynthesis in *C. crescentus,* perhaps contributing to the initiation or regulation of pilus formation.

### Deletion of *cpaL* increases surface adhesion and holdfast production

It has previously been demonstrated that pili play a crucial role in surface-sensing and adhesion in *C. crescentus* by rapidly stimulating synthesis of the holdfast upon surface-contact (11, 20). Therefore, we expected that the Δ*cpaL* mutant would exhibit a surface attachment defect due to reduced pilus synthesis. To test this hypothesis, we used a surface attachment assay (20) wherein cells were spotted onto a glass coverslip and incubated for 30 min to allow attachment to the surface. The coverslip was then washed to remove unattached cells, and the surface-adherent cells were imaged by microscopy. In this assay, holdfast-deficient cells do not attach, while the Δ*pilA* mutant exhibits around 54% reduction in surface binding compared to the *pilA-cys* holdfast positive (HF+) strain (Figure 3A). In contrast, the Δ*cpaL* mutant exhibited a three-fold increase in surface binding compared to the *pilA-cys* HF+ strain (Figure 3A, S4).

**Figure 3:**
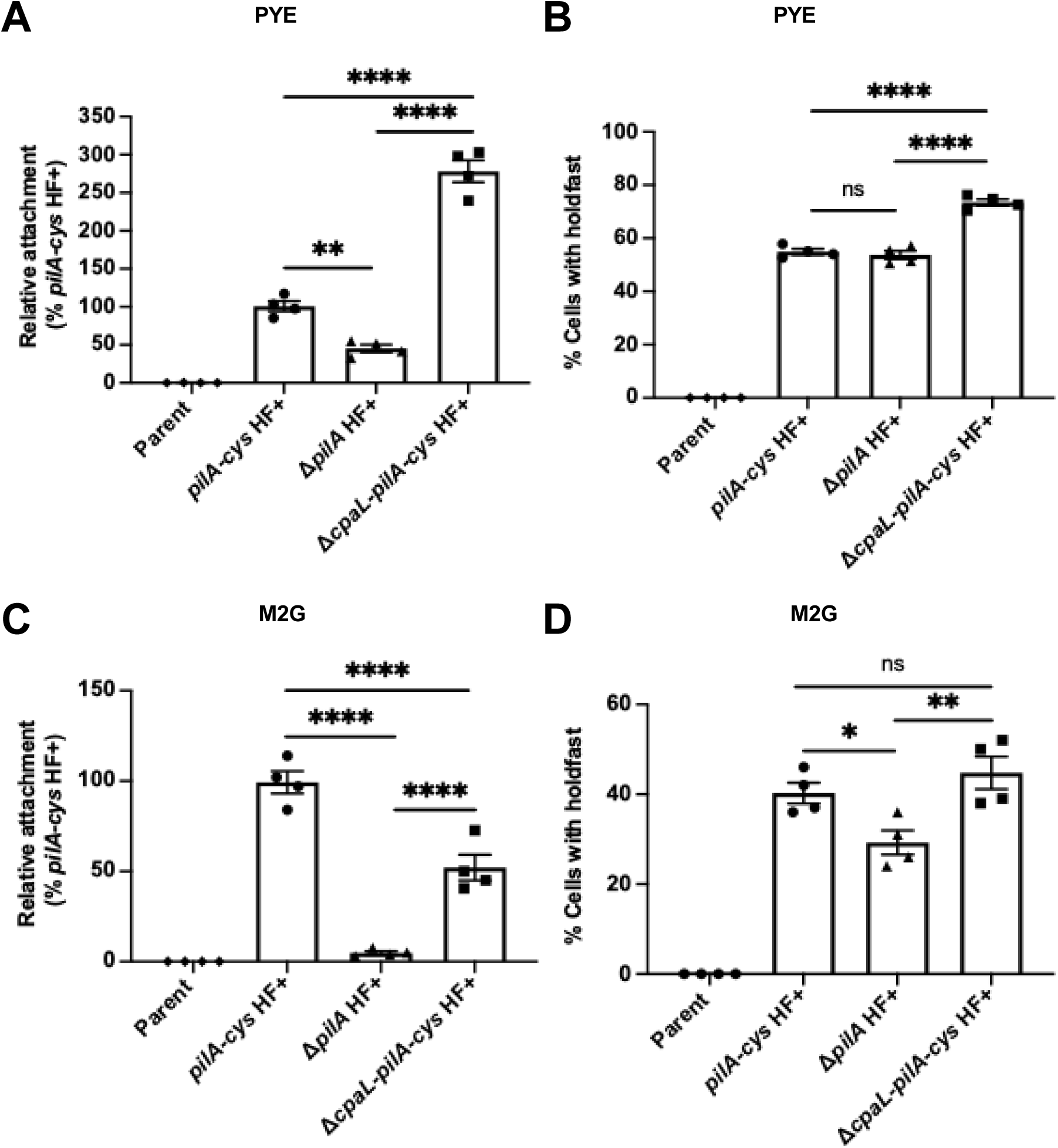
The deletion of *cpaL* increases surface attachment and holdfast production in PYE but not in M2G medium. **A:** Quantification of the attachment of cells grown in the complex medium PYE to a glass coverslip after 30 min of incubation relative to the strain *pilA*-cys HF+ (NA1000 *pilA*-cys, HF+: Holdfast positive). **B:** Quantification of the percentage of cells producing holdfast in the population in the complex medium PYE on an agarose pad. **C:** Quantification of the attachment of cells grown in the minimal medium M2G to a glass coverslip after 30 min of incubation relative to the strain *pilA*-cys HF+ (NA1000 *pilA*-cys, HF+: Holdfast positive). **D:** Quantification of the percentage of cells producing holdfast in the population in the minimal medium M2G on an agarose pad. Data are the mean of four independent biological replicates. Error bars indicate the standard error of the mean (SEM). Statistical comparisons were made using Tukey’s multiple comparisons test. ****, P <0.0001; **, P <0.01; *, P <0.05.

To determine whether the elevated surface adherence of the Δ*cpaL* mutant was due to increased holdfast production, we imaged holdfasts on agarose pads using a fluorescently labeled wheat-germ agglutinin lectin (AF488-WGA) that binds specifically to the *N*-acetyl-glucosamine moiety present in holdfasts (11, 24). We imaged on soft PYE agarose pads since they minimize the stimulation of holdfast production via the surface-sensing pathway (19, 25). Holdfasts were detected in 55% of the cells in the Δ*pilA* mutant and in the *pilA-cys* HF+ strain (Figure 3B). In contrast, 74% of the cells in the Δ*cpaL* mutant population had a holdfast (Figure 3B). Surface binding and holdfast production were restored to the levels of the *pilA-cys* HF+ strain when the Δ*cpaL* mutant was complemented with *cpaL* (Figure S5A and B). These results suggest that the increased surface adherence of the Δ*cpaL* mutant is attributable to elevated holdfast production, suggesting a role for CpaL in either the developmental or surface-sensing pathway of holdfast production.

### CpaL plays a role in surface-sensing

The Δ*cpaL* mutant exhibits increased surface adhesion and holdfast synthesis in the complex medium PYE (Figure 3A and B). In PYE, *C. crescentus* holdfast synthesis is regulated by both the developmental pathway, where holdfast is produced during differentiation from swarmer to stalked cells, and the surface-sensing pathway, where holdfast production is rapidly stimulated upon surface contact, hastening cell differentiation. In contrast, *C. crescentus* cells grown in the defined medium M2G do not undergo surface-contact mediated holdfast synthesis (19).

To determine whether the increased surface attachment and holdfast synthesis phenotypes of the Δ*cpaL* mutant are related to the developmental or surface sensing pathway of holdfast synthesis, we performed surface attachment and holdfast quantification assays in defined M2G medium, using identical conditions as in PYE (19). First, we found that pilus production in the Δ*cpaL* mutant was comparable between M2G and PYE, with a low proportion of cells synthesizing a single long pilus, a defect that was complemented by a plasmid copy of *cpaL* (Figure S3). However, in M2G medium, the Δ*cpaL* mutant exhibited a 48% reduction in surface attachment compared to the *pilA-cys* HF+ strain, as expected for a mutant with reduced pilus production (Figure 3C). Furthermore, approximately 42% of both the parental and Δ*cpaL* mutant populations produced holdfast, with no significant difference detected between these strains (Figure 3D). These results are in stark contrast to the increased surface adhesion and holdfast synthesis phenotypes of the Δ*cpaL* mutant seen in PYE. Since surface contact does not stimulate holdfast production in M2G, these results suggest that increased holdfast synthesis and surface attachment of the Δ*cpaL* mutant in PYE results from a role of CpaL in pilus-mediated surface-sensing.

To further examine the interaction of cells with a surface, we monitored the timing of holdfast synthesis in single cells in response to surface contact in PYE medium using a PDMS microfluidic device, where cells were introduced into a well of the device and allowed to adhere to a glass cover slip in the presence of AF488-WGA. The AF488-WGA allowed us to track the formation of holdfast in individual cells upon surface contact. While the *pilA-*cys HF+ strain showed rapid holdfast synthesis upon surface contact, within an average of approximately 26.13 ± 2.88 sec, the *ΔcpaL* mutant was comparatively slower in producing holdfast upon surface contact (106.4 ± 17.45 sec, Figure 4 A and B). The median holdfast synthesis times for the *pilA-cys* HF+ mutant and the Δ*cpaL-pilA-cys* HF+ mutant were 10 sec and 20 sec, respectively. These results indicate that *cpaL* plays a role in regulating holdfast production through the surface sensing pathway.

**Figure 4:**
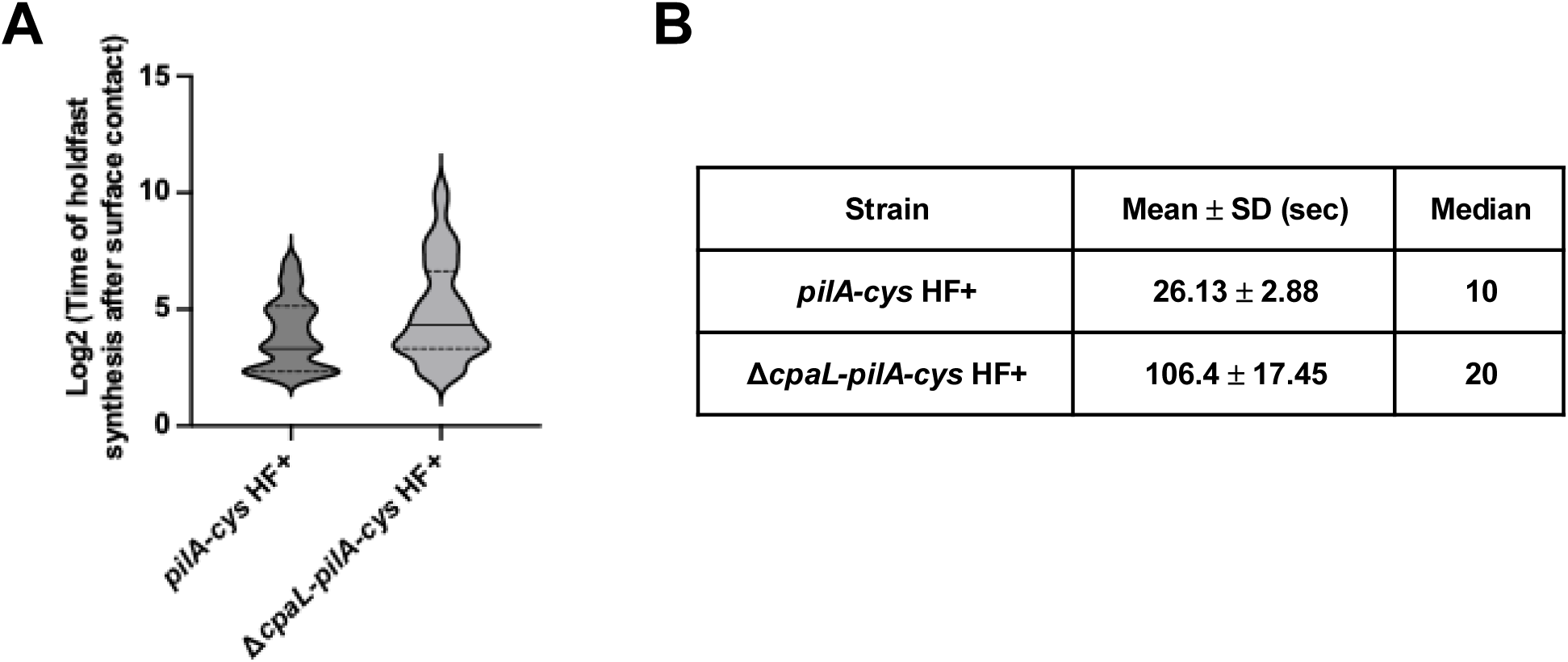
The absence of CpaL leads to a delay in holdfast production after surface contact. **A:** Logarithmic violin plot showing the time of holdfast synthesis after surface contact (sec), for there independent replicates of *pilA-cys* HF+ (n=119) and Δ*cpaL-pilA-cys* HF+ (n=163). Statistical comparison was made using a two-tailed unpaired T-test. ****, P <0.0001. **B:** Table showing the mean and the median of the strains used in the violin plot.

### The obstruction of pilus retraction does not affect surface attachment and holdfast production in the Δ*cpaL* mutant

The above results suggest that the deletion of *cpaL* elevates holdfast production through stimulation of the surface sensing pathway. To test if surface sensing could be further stimulated in the Δ*cpaL* mutant, we blocked pilus retraction using PEG5000-mal. This treatment has previously been shown to stimulate the production of holdfast via the surface sensing pathway, even in the absence of a surface (11). Specifically, we compared surface attachment and holdfast production among the different strains with and without PEG5000-mal to block pilus retraction. Blocking pilus retraction in the *pilA-cys* HF+ strain caused a strong increase in the percentage of surface-attached cells compared to the HF+ strain lacking the *pilA*-cys mutation and to the unblocked *pilA-*cys HF+ negative control (Figure 5A), consistent with previous results (11). In contrast, while the Δ*cpaL pilA-cys* mutant exhibited greater surface attachment compared to *pilA-cys* HF+ (Figure 3A), attachment of this mutant could not be further stimulated by PEG5000-mal (Figure 5C).

**Figure 5:**
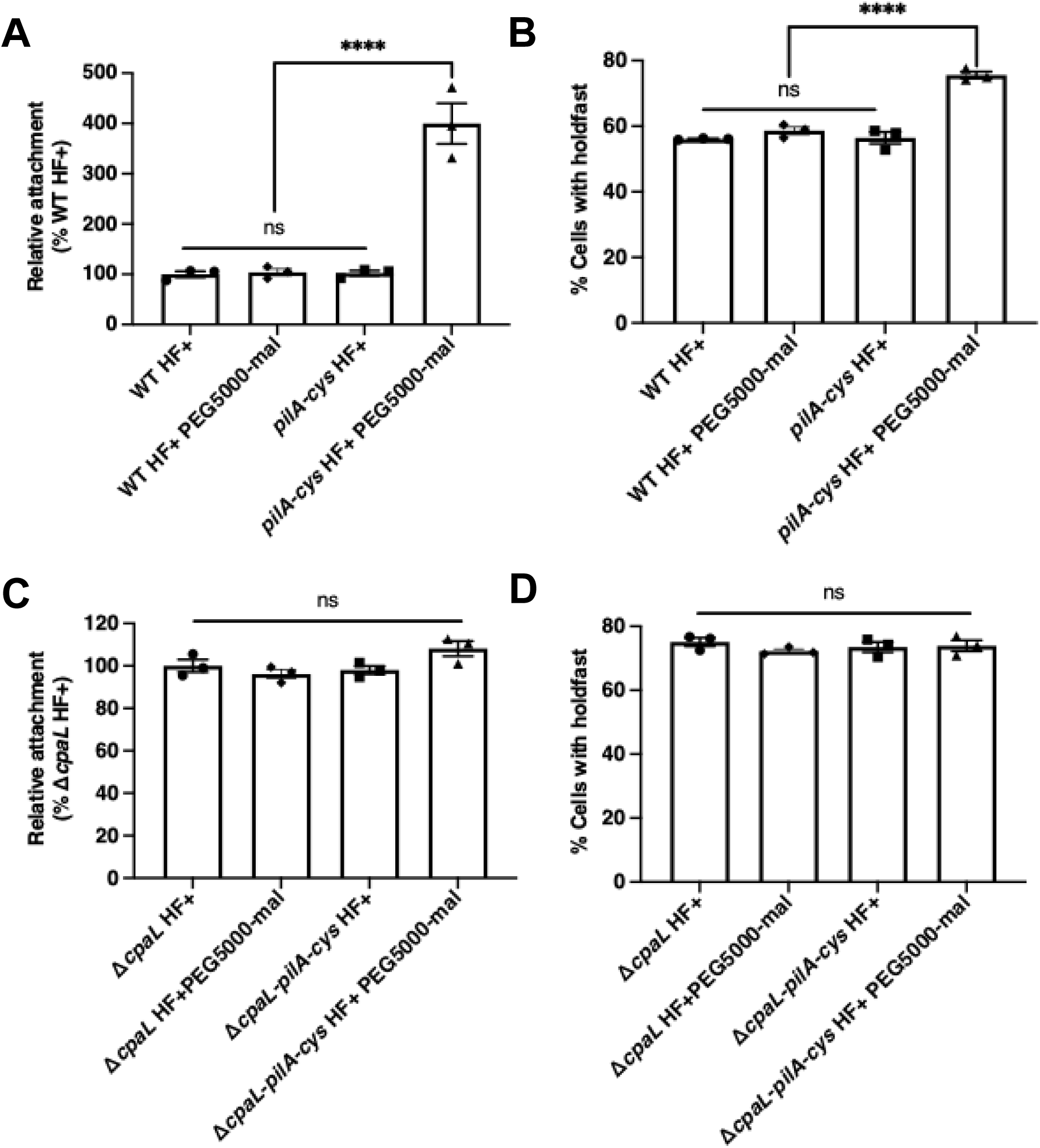
The Δ*cpaL* mutant is insensitive to the addition of PEG5000-mal: **A:** Quantification of the attachment of cells to a glass coverslip after 30 min of incubation relative to the strain WT HF+ (NA1000, HF+: Holdfast positive). PEG5000-mal was added to cultures 5 min before cells were added to the coverslip to block pili retraction in *pilA-cys* strains. **B:** Quantification of the percentage of cells producing holdfast in the population after 5 min of incubation with PEG5000-mal on agarose pad. The percentage of cells producing holdfast in the population was quantified after labeling holdfast with the AF488 conjugated wheat germ agglutinin (AF488-WGA). **C:** Quantification of the attachment of cells to a glass coverslip after 30 min of incubation relative to the strain Δ*cpaL* HF+ (NA1000, HF+: Holdfast positive). PEG5000-mal was added to cultures 5 min before cells were added to the coverslip to block pili retraction in *pilA-cys* strains. **D:** Quantification of the percentage of cells producing holdfast in the population after 5 min of incubation with PEG5000-mal on agarose pad. The percentage of cells producing holdfast in the population was quantified after labeling holdfast with the AF488 conjugated wheat germ agglutinin (AF488-WGA). Data are the mean of three independent biological replicates. Error bars indicate the standard error of the mean. Statistical comparisons were made using Tukey’s multiple comparisons test. ****, P <0.0001.

We next measured holdfast production in the presence of PEG5000-mal. Holdfast production was stimulated by PEG5000-mal in the *pilA-cys* HF+ strain compared to the HF+ strain without the *pilA-cys* mutation and the *pilA-cys* HF+ mutant in the absence of PEG5000-mal (Figure 5B), consistent with previous results (11). However, blocking pilus retraction in the Δ*cpaL pilA-cys* HF+ mutant did not further increase holdfast production compared to the untreated condition, or compared to the Δ*cpaL* HF+ mutant without the *pilA-cys* mutation, in either the blocked or unblocked conditions (Figure 5D).

Collectively, our results indicate that the absence of *cpaL* leads to a constitutive stimulation of the surface-sensing pathway and suggest that the *cpaL* gene plays a role not only in pilus biogenesis but also in some aspect of the surface-sensing mechanism.

### CpaL is predicted to adopt a pilin-like module linked to a vWA domain

To better understand the function of CpaL, we performed several sequence-based bioinformatic analyses to identify conserved domains and sequence features of CpaL. The predicted 3D structure of CpaL was generated using the AlphaFold3 server (26) (Figure 6A, S6A and B). The predicted structure was then submitted to the Dali server (27) to identify experimentally determined protein structures in the Protein Data Bank (PDB) that exhibit structural similarity to the predicted CpaL model (Figure 6A). The top structural analogs of CpaL determined by Dali are all von Willebrand factor type A (vWA)-domain containing proteins that play a role in pilus or fimbrial synthesis in bacteria (Table S4). The vWA domain is a widely distributed structural motif that, in bacteria, is characterized by its ability to mediate surface adhesion, adherence to host-derived proteins (28, 29), and mechanosensing (6) in conjunction with T4P.

**Figure 6:**
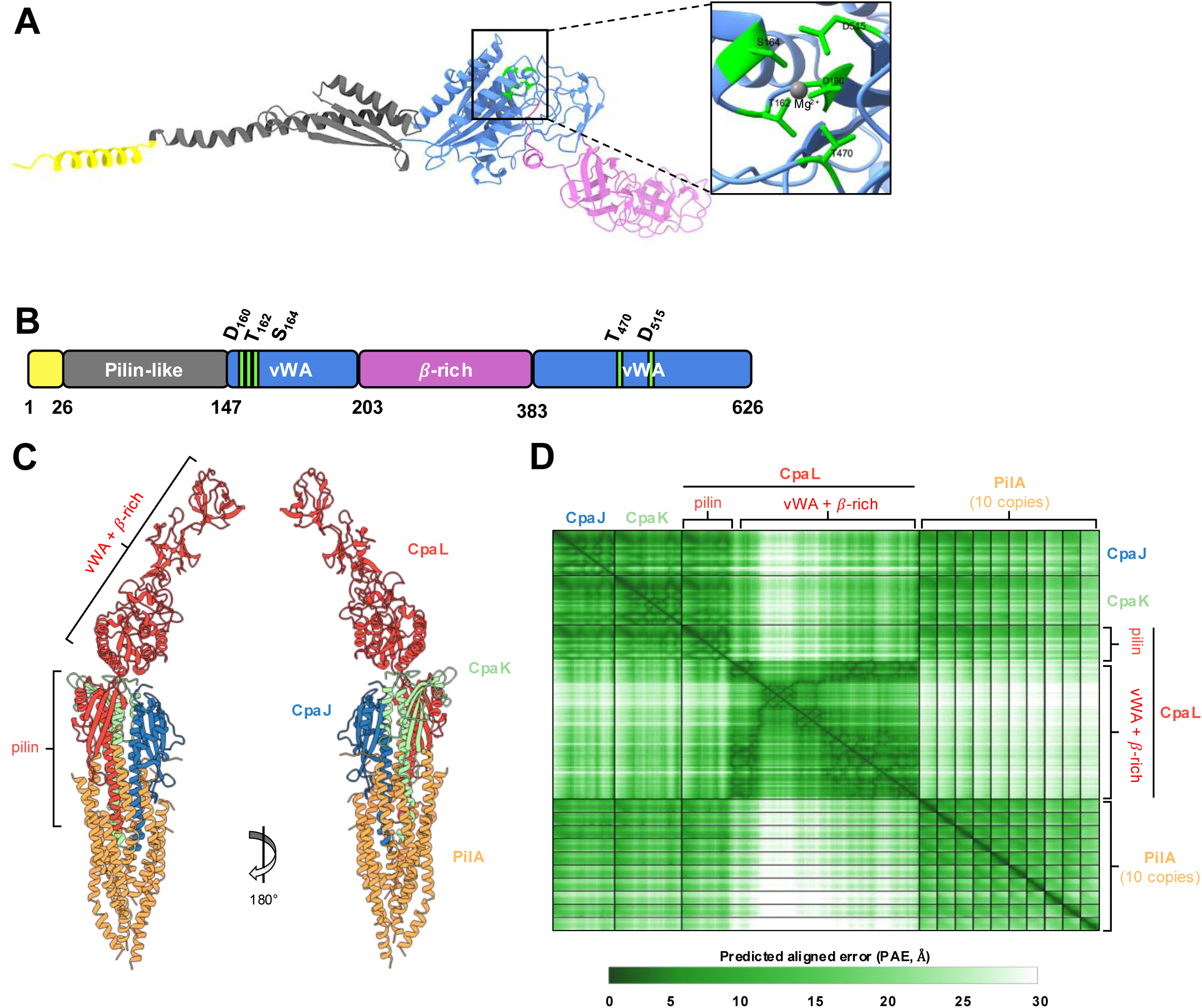
CpaL is predicted to contain a pilin-like module and a vWA domain. **A:** Ribbon diagram showing the predicted structure of CpaL generated using AlphaFold3. Domains and coloring are as described in panel B. The inset depicts the MIDAS motif in the vWA-like domain which is composed of five residues (D160, T162, S164, T470, and D515, green) with a predicted Mg^2+^ ion (gray) incorporated among those residues. **B:** Schematic representation of the CpaL domain organization. The first region at the N-terminus is predicted to be a signal peptide (residues 1-26, yellow), followed by a pilin-like module (residues 27-147, dark grey). A single von Willebrand Factor A-like domain (vWA) is divided between two distinct segments of the CpaL sequence (residues 148-203 and 384-626, blue), between which are two tandem β-rich domains (residues 204-383, pink). **C:** Top-ranked structure of a complex composed of the minor pilins CpaJ (blue), CpaK (green), CpaL (red), and ten copies of the major pilin subunit PilA (orange) predicted by AlphaFold3. The predicted or known signal sequences of each protein were removed prior to the prediction. **D:** The predicted aligned error (PAE) scores for the model depicted in panel C. The PAE indicates the positional error in angstroms (Å) for a given pair of residues across all protein chains in the model.

The N-terminus of CpaL contains a domain of approximately 120 residues predicted to adopt a pilin-like fold (residues 27-147): a long α-helix followed by a β-sheet consisting of three β-strands (Figure 6A and B, dark grey), giving this module its characteristic lollipop shape (16). The predicted pilin-like module of CpaL shares structural similarity with the T4P pilin THHA1221 from *Thermus thermophilus* (PDB: 4BHR) and the minor pilin PilX from *Neisseria meningitidis* (PDB: 2OPD), as determined by Dali, as well as the predicted structures of the *C. crescentus* Tad minor pilins CpaK and CpaJ (Figure S9B). Sequence analysis of this module revealed a potential cleavage site for the prepilin peptidase CpaA, which consists of a stretch of hydrophilic residues followed by the consensus sequence G/A - X_4_ - F/E, ending with a stretch of predominantly hydrophobic residues (16, 28) (Figure S9A).

The vWA-like domain is present at the C-terminus of CpaL and is formed by two sections of the polypeptide chain separated by a β-rich domain (residues 148-203 and 384-626) (Figure 6A and B, blue). The vWA domain of CpaL is predicted to adopt a classical Rossmann fold that is broadly conserved across all vWA-like domains (30). Alignment of the vWA domain of CpaL with the vWA domain of the top Dali hit, SpaC of *L. rhamnosus* GG (PDB ID: 6M48) (Table S4), revealed significant structural similarity in the core Rossmann fold of the vWA domains of these proteins, with a 1.039 Å root mean square deviation (RMSD) across 118 atom pairs. The vWA domain of CpaL also contains a sequence motif known as the metal ion-dependant adhesion-site (MIDAS) motif (Figure 6A and B, S10A and B), which is often present in vWA domains, where it coordinates a divalent metal ion (31). This motif is characterized by the DxS/TxS/T, T, and D signature (32), which in CpaL corresponds to D_160_, T_162_, S_164_, T_470_, and D_515_, respectively, located at the top of the central β-sheet, where they are predicted to coordinate an Mg^2+^ ion (Figure 6A and B, S10A and B). The primary sequence that comprises the CpaL vWA domain flanks an intervening β-rich region (residues 204-384, purple) that is projected away from the MIDAS motif and contains two consecutive six-stranded β-barrels (Figure 6A and B). A Dali search using only the structure of the β-rich region did not return any hits with strong similarity to this region of CpaL.

### CpaL is predicted to assemble at the pilus tip

The presence of a pilin-like domain in the predicted structure of CpaL (Figure 6A and S9B) and the similarity of this structure to pilus tip-localized minor pilins (Table S4) suggests that CpaL may function in the context of the pilus tip. To examine this, we utilized AlpahFold3 (26) to predict a putative Tad pilus tip complex composed of CpaL, the minor pilins CpaJ and CpaK, and ten copies of the major pilin subunit PilA to simulate a pilus filament. The sequences of the predicted or known signal sequences of each of these proteins were removed prior to the prediction. This yielded a structure in which CpaJ, CpaK, and the pilin-like domain of CpaL form a trimeric complex (Figure 6C, S7) that sits atop the ten PilA subunits (Figure 6C, S8A and B), which are themselves arranged similarly to the recently resolved structure of the PilA filament (Figure S8C; PDB 8U1K) (33). The transmembrane helices of CpaJ, CpaK, and CpaL are incorporated into the distal end of the filament-like PilA structure, while the vWA domain of CpaL projects away from the rest of the complex on the opposite side (Figure 6C, S8A and B). Importantly, predicted aligned error (PAE) scores for the model, particularly in the region comprising the interaction between CpaJ, CpaK, and the pilin-like domain of CpaL, are low (Figure 6D, S7C and S8A), suggesting that this could represent a biological arrangement of these proteins.

### CpaL functions differently than the other minor pilins CpaJ and CpaK

Because of the similarity of CpaL’s pilin-like domain to the minor pilins CpaJ and CpaK (Figure S9B) and its predicted assembly with these minor pilins to form a complex at the tip of the pilus (Figure 6C and D, S7 and S8), we sought to determine if the minor pilins CpaJ and CpaK play a similar role in the surface sensing pathway. We tested the effect of *ΔcpaJ* and *ΔcpaK* deletions on surface adhesion and holdfast synthesis and found that deletion of these two minor pilin genes phenocopied *ΔpilA* cells – they were resistant to the pilus-dependent phage ΦCbK (Figure 7A), lacked pili (Figure 7B), had significantly reduced surface adhesion (Figure 7C), and produced holdfasts similarly to WT (Figure 7D). Thus, we conclude that the function of CpaL is different from the function of the minor pilins CpaJ and CpaK.

**Figure 7:**
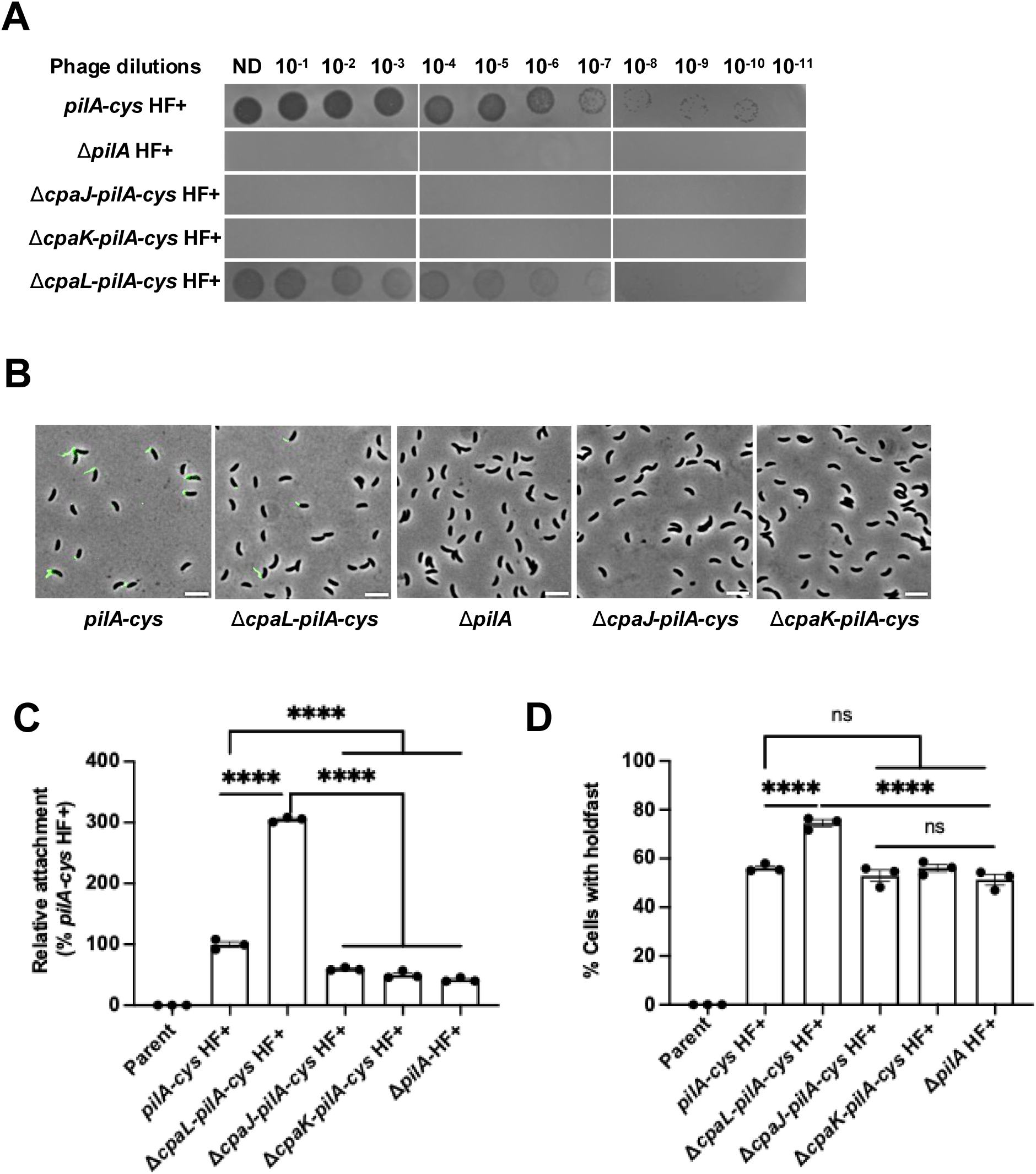
The deletion of minor pilin genes *cpaJ* and *cpaK* phenocopies *ΔpilA*. **A:** Top agar phage sensitivity assays. Serial dilutions of the phage were spotted on plates with top agar containing each bacterial strain. ND, no dilution. **B:** Representative microscopy images of synchronized swarmer cells that were blocked for pilus retraction with PEG5000-mal and labeled with AF488-maleimide (green). Scale Bar, 10 μm. **C:** Quantification of the attachment of cells grown in the complex medium PYE to a glass coverslip after 30 min of incubation relative to the strain *pilA*-cys HF+ (NA1000 *pilA*-cys, HF+: Holdfast positive, Parent: NA1000 HF-). **D:** Quantification of the percentage of cells producing holdfast in the population in the complex medium PYE on an agarose pad. Data are the mean of three independent biological replicates. Error bars indicate the standard error of the mean (SEM). Statistical comparisons were made using Tukey’s multiple comparisons test. ****, P <0.0001.

In summary, CpaL is predicted to adopt an N-terminal pilin-like fold that may allow it to incorporate into the tip of the pilus filament alongside the minor pilins CpaJ and CpaK, while its C-terminal vWA domain may play a role in adhesion and/or surface-sensing.

## Discussion

In this study, we demonstrate the importance of CpaL for pilus biosynthesis in *C. crescentus* and show that the absence of CpaL influences holdfast production, likely through stimulation of the surface-sensing pathway. In *C. crescentus,* the absence of pili reduces adherence to a solid substratum (11). However, while deletion of *cpaL* dramatically reduces pilus production (Figure 2), this mutation leads to an unexpected increase in surface attachment and holdfast synthesis (Figure 3A and B). These results prompted us to explore whether CpaL may perform a role in pilus-mediated surface sensing. To examine this, we cultured *C. crescentus* in M2G minimal medium, which prevents holdfast synthesis via the surface sensing pathway but still allows for developmentally-regulated holdfast synthesis (19). Under these conditions, we found that the Δ*cpaL* mutant exhibits reduced attachment to glass, as would be expected for a mutant with impaired pilus biosynthesis (Figure 3C), and no longer produces increased levels of holdfast (Figure 3D). Together, these results suggest that deletion of *cpaL* stimulates holdfast biosynthesis and attachment through the surface-sensing pathway. This hypothesis is supported by the absence of further stimulation of holdfast production in the Δ*cpaL* mutant after artificially blocking pilus retraction with PEG5000-mal, which strongly boosts holdfast production and attachment in the *pilA-cys* HF+ strain (Figure 4A and B). Thus, it appears that the surface-sensing pathway is stimulated constitutively in the Δ*cpaL* mutant, suggesting that CpaL plays a role in this pathway.

To gain insight into how CpaL may be performing seemingly opposing roles in pilus production and surface sensing, we examined the predicted structure of CpaL using Alphafold3. Analysis of its structure identified an N-terminal pilin-like module with a structural arrangement that is characteristic of pilin proteins (16) (Figure 6A, B, S9A, and B), with particular structural homology to the major pilin THHA1221 from *T. thermophilus* (PDB: 4BHR) (34) and the minor pilin PilX from *N. meningitidis* (PDB: 2OPD) (35) (Figure S9B). The pilin-like module of CpaL also shares similarity with the predicted structures of the *C. crescentus* minor pilins, CpaJ and CpaK (Figure S9B). A potential cleavage site for the pre-pilin peptidase CpaA was identified in the pilin domain of CpaL (Figure S9A), which suggests that CpaL could be processed to a mature form to facilitate incorporation into the pilus fiber, as occurs for *C. crescentus* PilA, and potentially also for CpaJ and CpaK (Figure S9A). Position +5 after the peptidase cleavage site in T4aP and T4bP pilins is a highly conserved Glu that forms a salt bridge with the N-terminal amine of the previously-incorporated pilin subunit, which is critical for fibre assembly. Exceptions to this rule are noted for a subset of minor pilins, the GspK orthologs, which instead have a non-polar residue at position +5. It is hypothesized that the absence of the conserved Glu5 in these minor pilins is indicative that they are the first component to comprise the nascent filament(16). However, archaeal pilins lack the conserved Glu5, and although this residue is present in bacterial Tad pilins, the structure of the *C. crescentus* PilA filament demonstrated that Glu5 points towards the solvent and does not participate in intermolecular interactions (33). Instead, the adjacent Tyr6 is oriented below the N-terminal Ala1 of the previously-incorporated pilin subunit, maintaining the helical register of the filament. Therefore, Tyr6 in Tad pilins may fulfill a parallel function to Glu5 in T4aP and T4bP pilins. However, CpaL possesses an Ala at position +6, which may indicate that, like the GspK orthologs, CpaL is the first subunit to comprise the nascent filament. Indeed, in our AlphaFold3-predicted structure of CpaL at the pilus tip (Figure 6C), CpaL is oriented as the first molecule in the filament. These predicted structural features collectively suggest that CpaL could be a Tad minor pilin in *C. crescentus*, albeit of unusual size and architecture. Indeed, the CpaL ortholog from the Tad pilus system of *actinomycetemcomitans*, TadG, has previously been identified as a constituent of pilus fiber preparations (36–38).

Minor pilins with atypical domain architectures have recently been structurally resolved in the T4P system of *S. sanguinis* (28, 39), where the minor pilins PilB and PilC were found to have a vWA domain and a glycan binding domain, respectively, linked to a pilin module. These studies have further demonstrated that the tripartite minor pilin complex of *S. sanguinis*, composed of PilA, PilB, and PilC, forms an “open wings” architecture that can only be accommodated at the tip of the pilus fiber (39, 40). The incorporation of such a minor pilin complex at the pilus tip is consistent with what is observed in the T4aP systems of *P. aeruginosa* and *M. xanthus*. In these species, an inner membrane complex composed of several minor pilins, as well as the vWA domain containing non-pilin subunit PilY1, recruits major pilin subunits to initiate pilus fiber assembly. This minor pilin-PilY1 complex is ultimately incorporated into the tip of the growing pilus fiber after successful nucleation of pilus assembly (41, 42). Thus, CpaL, together with the putative Tad minor pilins CpaJ and CpaK, could similarly form a complex that is incorporated at the tip of the pilus fiber in *C. crescentus*. Indeed, our structural predictions with AlphaFold3 suggest that CpaL forms a tripartite complex with CpaJ and CpaK (Figure S7 and S8) that could be accommodated at the distal end of the pilus filament (Figure 6C and S8). Based on this hypothesis, it is possible that the reduced pilus production phenotype of the Δ*cpaL* mutant is due to disruption of this complex, resulting in fewer pilus nucleation events. Infrequent nucleation could also explain why the pili of the Δ*cpaL* mutant are longer on average: infrequent pilus nucleation events would result in a larger pool of major pilin subunits in the inner membrane, drawn from to assemble fewer pili (typically one per piliated cell). However, CpaL does not simply play the role of a minor pilin since deletion of *cpaJ* or *cpaK* yields cells without pili and without the stimulation of surface adhesion and holdfast synthesis seen for *ΔcpaL*.

In addition to a pilin-like module, CpaL also has a predicted vWA domain (6, 30, 43) (Figure 6A, B, S6A, B, and S11). T4P-associated proteins in Gram-positive and Gram-negative bacteria harboring vWA domains are often directly involved in surface adhesion (9, 28, 29). Many vWA domains contain a five-residue motif called the MIDAS motif, which is often required to mediate surface adherence (28). For example, the MIDAS motif from the T4P associated protein PilC1 in *Kingella kingae* is required for adherence to host tissues as well as for twitching motility (44). CpaL is also predicted to contain a canonical MIDAS motif that coordinate a metal ion Mg^2+^ (Figure 6A, B, and S11A, B), supporting CpaL’s role in surface adhesion. Additionally, it is well-documented that vWA domain containing proteins have a mechanosensory function in eukaryotes (30). Thus, it is thought that vWA domains could also play a role in T4P-mediated mechanosensing in bacteria (6, 9). Indeed, in the pilus-tip associated protein PilY1 in *P. aeruginosa,* the vWA domain undergoes sustained conformational changes when force is applied by AFM. This mechanosensitivity is perturbed by the mutation of a critical disulphide-bond in the protein, which also reduces surface-sensing behaviours in *P. aeruginosa*, suggesting a link between force-induced conformational changes in PilY1 and surface sensing (6, 9). Given the CpaL’s role in surface sensing, its predicted structure with a vWA domain, and its hypothesized localization to the pilus tip, a similar mechanism could therefore be at play in CpaL-mediated surface sensing in *C. crescentus*.

Despite the similarities of CpaL to PilB in *S. sanguinis* and PilY1 in *P. aeruginosa*, the increased surface attachment of the *C. crescentus* Δ*cpaL* mutant stands in contrast to what has been reported for the *pilB* and *pilY1* mutants, which exhibit decreased biofilm formation, consistent with their loss of pilus production (45, 46). In fact, loss of pili in *C. crescentus* through deletion of the major pilin *pilA* also reduces biofilm formation through abolishment of pilus-mediated surface-sensing (11). However, deletion of *cpaL* stimulates the production of holdfast via the surface-sensing pathway, similar to what is observed when pilus retraction is artificially blocked. It has previously been shown that binding of bulky PEG5000-mal adducts to the pilus fiber prevents pilus retraction by sterically occluding the entrance of the modified pilus filament into the CpaC secretin pore in the outer membrane, thereby stimulating the surface sensing pathway (20). Similarly, stimulation of surface sensing was observed when a glycine to aspartate mutation was introduced into CpaC in the predicted outer lip of the pilus secretin pore (20), again pointing to interactions between the pilus filament and the secretin as key to the mechanism of surface sensing. Based on our results, we hypothesize that CpaL mediates the surface sensing pathway by modulating how the pilus fiber interfaces with CpaC and rest of the Tad pilus secretion machinery. Evidence from *M. xanthus* and *P. aeruginosa* suggests that pilus tip proteins remain assembled as a priming complex through successive rounds of pilus extension and retraction, and that this complex acts as a plug in the secretin pore to prevent full retraction of the pilus into the inner membrane (47) (48). In these species, pilus retraction may position PilY1 within the secretin pore, where force-induced conformational changes in PilY1 due to surface contact (9) may hypothetically be sensed. By analogy, conformational changes in CpaL due to surface contact may similarly be sensed by CpaC to stimulate the surface sensing pathway in *C. crescentus*. However, in the absence of CpaL, the pilus tip may interact with the secretin in such a way that surface-sensing is constitutively triggered. Further investigation is required to better understand the exact role of CpaL in the surface-sensing mechanism.

In conclusion, our results demonstrate the importance of CpaL for pilus formation and for regulation of holdfast biosynthesis through the surface-sensing pathway. Structural predictions of CpaL reveal similarity to the mechanosensitive vWA domain and possible incorporation at the tip of the pilus fiber via a pilin-like module. Together, these results implicate CpaL as a candidate mechanosensor for the Tad pilus in *C. crescentus*.

## Material and Methods

### Bacterial strains, plasmids, and growth conditions

The bacterial strains used in this study are listed in Table S1. *Caulobacter crescentus* strains were grown at 30°C in peptone-yeast extract (PYE) (49) or in defined M2 medium supplemented with 0.2% (w/v) glucose (M2G) (50). PYE was supplemented with 5 μg/ml kanamycin (Kan), where appropriate. Commercial, chemically competent *Escherichia coli* DH5-α (Bioline and New England Biolabs) was used for plasmid construction and was grown at 37°C in lysogeny broth (LB) supplemented with 25 μg/ml Kan, where appropriate.

Plasmids (Table S2) were transferred to *C. crescentus* by electroporation, transduction with ΦCr30 phage lysates, or conjugation with S17-1 *E. coli* as described previously (51). In-frame deletion strains were made by double homologous recombination using pNPTS-derived plasmids as previously described (52). Briefly, plasmids were introduced into *C. crescentus*, and then two-step recombination was performed using kanamycin resistance to select for single crossover followed by sucrose resistance to identify plasmid excision events. All mutants were validated by sequencing to confirm the presence of the deletion.

### Plasmid construction

The *cpaL* complementation construct was made using the high-copy-number vector pBXMCS-2 (53) with *cpaL* under the control of a xylose-inducible promoter. Because expression from this promoter is leaky, no xylose was added to growth media for *cpaL* expression. The *cpaL* open reading frame was amplified from *C. crescentus* NA1000 genomic DNA using the cpaL-F and cpaL-R primers (Table S3). The *cpaL* PCR fragment was digested using NdeI and EcoRI and ligated into pBXMCS-2 digested by the same enzymes. Clones with a positive insert were verified by Sanger sequencing.

### Screen of gene for pilus retraction

500 µl of an overnight culture *of pilA-cys Caulobacter* cells were mixed with 50 µl of exponential *E. coli* carrying pFD1 (Rubin et al. PNAS 1999) and briefly vortexed. The mixture was pipetted onto a 0.22 µm filter on a vacuum manifold to concentrate cells to encourage conjugation, and the filter containing the cell concentrate was placed cell-side up on a plain PYE plate and incubated overnight at room temperature. Filter-concentrated cells were resuspended in 500 µl of PYE medium and mixed with 500 µl of undiluted ∼10^10^ pfu/ml ϕCbK phage stock (MOI of ∼10^6^) and incubated at room temperature for 10 min. 100 µl aliquots of cell/phage mixture were then plated on PYE plates containing kanamycin to select for transposon mutants and nalidixic acid to select against *E. coli* cells and grown at 30°C until colonies appeared (1-2 days). 184 colonies from kanamycin/nalidixic acid plates were inoculated into 96-well plate wells containing 200 µl of PYE and grown overnight at 30°C. Cells in wells were mixed with 20 µl of DMSO and stored at -80°C for long-term storage.

Individual mutants were then inoculated into 3 ml of PYE medium and grown overnight at 30°C. To assess pilus phenotypes, overnight cultures were labeled with 25 µg/ml AF488-mal and incubated at room temperature for 5 minutes. To remove excess dye, cultures were centrifuged at 7,500 *x g* for 1 min, the supernatant was removed, and cells were resuspended in 100 µl fresh PYE before imaging. 30 mutants exhibited labeled T4P and/or fluorescent cell bodies indicative of T4P production and were further characterized. Irradiated ϕCr30 lysates of isolated transposon mutants were generated and transduced into the parent strain YB8288 to confirm that phage resistance and T4P phenotypes were dependent on transposon insertions. Strains exhibiting complete or partial phage resistance along with T4P fluorescence phenotypes where further characterized. Genomic DNA was extracted from each strain and digested using the restriction enzyme Sau3AI at 37°C for 1 hr followed by heat inactivation at 65°C for 20 min. Digested DNA was then ligated in 100 µl reactions to promote self-annealing using T4 phage ligase at room temperature for 1 hr. Ligations were then used as templates for inverse PCR reactions using primers Mariner F and Mariner R targeting the transposon element and sequenced using the same primers. Sequences adjacent to the transposon element were mapped to the *Caulobacter crescentus* genome to identify transposon insertion sites.

### Phage sensitivity assays

Phage sensitivity assays were performed using the pilus-specific phage ΦCbK as described previously (23). Briefly, 200 µl of *C. crescentus* stationary phase culture was mixed with 3 ml of 0.5% (w/v) soft PYE agar. The mixture was spread over a 1.5% (w/v) PYE agar plate and incubated at room temperature for 1 h to solidify. A tenfold serial dilution series of ΦCbK was prepared in PYE, and 5 µl of each dilution was spotted on top of the agar plate. The plates were grown for 2 days at 30°C before imaging using a ChemiDoc MP (BioRad).

### Growth rate analysis

Growth of *C. crescentus* strains was measured in 24-well polystyrene plates (Falcon) using a SpectraMax iD3 microplate reader (Molecular Devices). Stationary-phase cultures were diluted to an optical density at 600 nm (OD_600_) of 0.05, and 1 ml of this culture was added to the wells of the microplate in triplicate and incubated for 24 h at 30°C under shaking. The OD_600_ was measured at 30 min intervals to generate growth curves (OD_600_ versus time).

### Synchronization of *C. crescentus* populations

*C. crescentus* populations were synchronized to enrich for pilus-producing swarmer cells, enabling facile quantification of pilus characteristics. The swarmer cells were synchronized and collected as described previously (54) with some modifications. To do this, *C. crescentus* cultures were grown to an OD_600_ of ∼ 0.15-0.3, and 2 ml of this culture was centrifuged at 5,400 × *g* for 5 min. The supernatant was removed, and the cell pellet was resuspended in 280 µl of PYE. Following this, 120 µl of polyvinylpyrrolidone colloidal silica solution (Percoll, Sigma) was added and mixed by gentle inversion. This mixture was centrifuged at 11,000 *× g* for 15 min, generating two bands of distinct cell populations: stalked cells in the upper band and swarmer cells in the lower band. 15 µl of the swarmer cell band was removed and added to 85 µl of PYE. This solution of synchronized swarmer cells was washed with 100 µl of PYE, before proceeding to pilus labeling, as described below.

### Pilin labeling, blocking, imaging, and quantification

The pili of *C. crescentus* were labelled as described previously (11). Briefly, 25 µg/ml of Alexa Fluor 488 C_5_ Maleimide (AF488-maleimide, ThermoFisher Scientific) was added to 100 µl of synchronized swarmer cell culture, prepared as described above, and incubated for 5 min at room temperature. Labelled cultures were centrifuged for 1 min at 5,400 *× g*, and the pellet was washed once with 100 µl of PYE to remove excess dye. The labelled cell pellets were resuspended in 7 to 10 µl of PYE, 0.5 µl of which was spotted onto a 1% agarose PYE pad (SeaKem LE, Lonza Bioscience). The agarose pad was sandwiched between glass coverslips for imaging.

To artificially block pilus retraction, methoxy-polyethylene glycol maleimide (PEG5000-maleimide, Sigma) with an average molecular weight of 5 kDa was used. 500 μM PEG5000-mal was added to 100 µl of synchronized swarmer cell cultures immediately prior to the addition of 25 μg/ml AF488-mal. Cultures were further prepared and imaged as described above.

For observing pilus dynamics, *C. crescentus* cells were grown to an OD_600_ of ∼ 0.15-0.3, labelled with AF488-maleimide as mentioned above, and 0.5 µl of the labelled sample was spotted onto a 1% agarose PYE or M2G medium pad as appropriate (SeaKem LE, Lonza Bioscience). The agarose pad was sandwiched between glass coverslips for imaging. Pilus dynamics were detected by time-lapse microscopy every 3 sec for 1 min.

Imaging was performed using a Nikon Ti-E inverted fluorescence microscope with Plan Apo 60X or 100X objectives, a GFP filter cube, an Andor iXon3 DU885 EM CCD camera, and Nikon NIS Elements imaging software.

The number of pili per cell, the percentage of piliated cells in the whole cell population, and the percentage of cells with fluorescent cell bodies were quantified manually using ImageJ software (55).

### Surface binding assay and holdfast quantification

Surface binding assays and holdfast quantification were carried out in parallel on the same samples. For each strain to be analyzed, a single colony was isolated and used to inoculate 3 ml of PYE or M2G medium, as appropriate. Tenfold serial dilutions of this culture (10^-1^ to 10^-4^) were prepared and incubated overnight at 30°C. The OD_600_ of each culture was measured, and the culture with an OD_600_ of ∼ 0.05 was selected. Different cultures with an OD_600_ of ∼ 0.05 were normalised exactly to OD_600_ = 0.05 before proceeding.

For each culture prepared as described above, 15 µl was transferred onto a glass coverslip and incubated in dark and humid conditions (to prevent desiccation and effects from varying light levels) for 30 min at 30°C. After incubation, coverslips were extensively washed in water, and a 1% PYE or M2G agarose pad, as appropriate, was added on top of the cells on the coverslip for microscopy analysis.

To quantify holdfast production, cells from the same culture preparations described above (OD_600_ = 0.05) were labelled with 0.5 µg/ml AF488-WGA (Wheat germ agglutinin lectin, ThermoFisher Scientific) for 1 min before spotting them onto a 1% agarose PYE or M2G pad, as appropriate, for microscopy analysis. WGA binds specifically to the N-acetylglucosamine residues present in the holdfast polysaccharide (24).

Cells attached to the coverslip, and labelled holdfast, were imaged using a Nikon Ti-E fluorescence microscope as described above. The percentage of cells attached to the coverslip, and the percentage of cells with holdfast, were quantified manually from microscopy images using ImageJ software (55).

To quantify surface binding and holdfast production from cells with blocked pili, 500 μM PEG5000-mal was added to cell cultures and incubated for 5 min before performing the experiments described above.

### Timing of holdfast synthesis after surface contact

Microfluidic well devices were constructed from PDMS (Polydimethylsiloxane) as described previously (11, 25). Briefly, a 10:1 mixture of PDMS prepolymer:curing agent was poured into a sterile petri plate, then the plate was placed for a few hours under a vacuum until the bubbles were removed. The plates were placed for 3h in an oven at 65°C. A rectangle of 25 x 35 mm was cut from the PDMS plates, with fluid access holes of 3 mm in diameter punched at 5 mm intervals. Next, the glass coverslip and the PDMS cast were assembled after plasma treatment and put overnight in an oven at 65°C.

For sample preparation, 200 ml of bacterial cell culture of OD_600_ = 0.6-0.8 grown in PYE was diluted in 800 ml of PYE containing 0.5 mg/ml AF488-WGA (Wheat germ agglutinin lectin, ThermoFisher Scientific) for holdfast labeling, then 10-15 μl of the above mixture was added to one well of the PDMS device.

Time-lapse videos were taken every 5 sec for 30 min at the glass-liquid interface using a Nikon Ti-E fluorescence microscope with Apo 60X objective, a GFP filter cube, an Andor iXon3 DU885 EM CCD camera, and Nikon NIS Elements imaging software. The attaching cells were detected using the phase contrast channel while holdfast synthesis was detected using the GFP channel. The time difference between holdfast synthesis and cell-surface attachment was determined using ImageJ software (55).

### CpaL structural and functional predictions

Prediction of protein domains, their global distribution, and associated architectures was done using the Dali (27) server which was used for comparing protein structures in 3D after generation of the predicted structure of CpaL using AlphaFold3 (26). The structure of CpaL in complex with CpaJ and CpaK at the pilus tip was predicted using AlphaFold3 (26). Molecular visualization of 3D protein structures was done using the ChimeraX software (56) and AlphaFold3 (26). The GenBank accession number for the primary sequence of CpaL is WP_010918088.1 and the UniProt Primary accession number is A0A0H3C404. The superposition of the vWA domain of CpaL with the vWA domain of SpaC (PDB ID: 6M48-A) was performed using ChimeraX software (56), while the superposition of the AlphaFold3-predicted PilA helical structure with the experimentally-determined structure of the pilus filament (PDB 8U1K), and calculation of alignment statistics, was done using MM-align (57).

## Supporting information

Movie S1 and will be used for the manuscript on the preprint site

Movie S2 and will be used for the manuscript on the preprint site

Movie S3 and will be used for the manuscript on the preprint site

Movie S4 and will be used for the manuscript on the preprint site

## Acknowledgment

This work was supported by a Fonds de Recherche du Québec, Nature et Technologies (FRQNT) doctoral training scholarship, a Bourse de Mérite from the Faculté de Médecine at the Université de Montréal, and a Bourse de formation à la recherche Mitacs to F.O.C.; a Natural Sciences and Engineering Research Council of Canada (NSERC) and FRQNT postdoctoral fellowship to G.B.W.; fellowship 1342962 from the National Science Foundation (NSF) to C.K.E.; and the Canada 150 Research Chair in Bacterial Cell Biology and grant R35GM122556 from the National Institutes of Health to Y.V.B. We also thank members of the Brun Lab for their feedback and critical review of the manuscript.

## Movies

**Movie S1:** Time-lapse movie of labeled non-synchronized parent cells showing pili extension-retraction dynamics after labeling with AF488-maleimide (green). Capture rate is 3 sec per frame. Frame rate is 5 fps. Scale Bar, 5 μm.

**Movie S2:** Time-lapse movies of labeled non-synchronized *cpaL* mutant cells showing pili extension-retraction dynamics after labeling with AF488-maleimide (green). Capture rate is 3 sec per frame. Frame rate is 5 fps. Scale Bar, 5 μm.

**Movie S3:** A representative time-lapse movie of cells producing holdfast upon surface contact for the parent in the presence of AF488-WGA. The cell bodies are in gray and the holdfasts are in green. Capture rate is 5 sec per frame. Frame rate is 25 fps. Scale Bar, 5 μm.

**Movie S4:** A representative time-lapse movie of cells producing holdfast upon surface contact for the *cpaL* mutant in the presence of AF488-WGA. The cell bodies are in gray and the holdfasts are in green. Capture rate is 5 sec per frame. Frame rate is 25 fps. Scale Bar, 5 μm.

---

**Figure S1:**
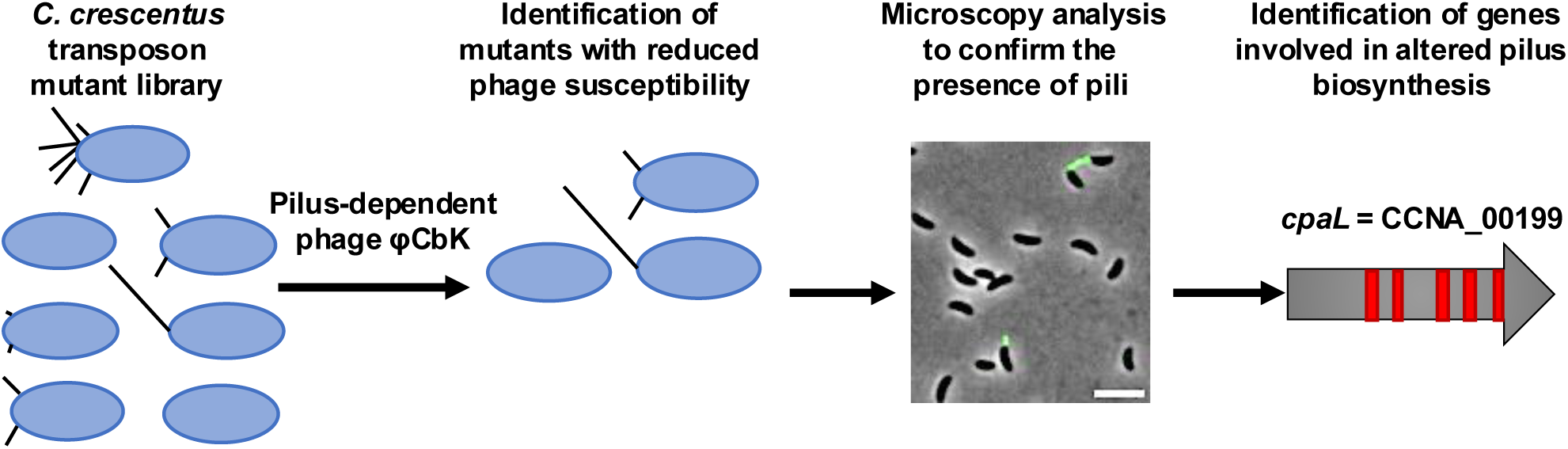
Genetic screen for resistance to a pilus-dependent phage identifies *cpaL* as a mediator of pilus activity. Schematic of the forward genetic screen used to identify mutants that are resistant to the pilus-dependent *Caulobacter* phage, ɸCbK, despite having pili. A *C. crescentus Mariner* transposon mutant library was generated and mixed with ɸCbK and grown on plates. Phage resistant mutants were isolated and imaged for pilus synthesis. Red bars indicate the position of transposon insertions identified within the *cpaL* (CCNA_00199) gene coding sequence (gray arrow). Scale bar, 10 μm.

**Figure S2:**
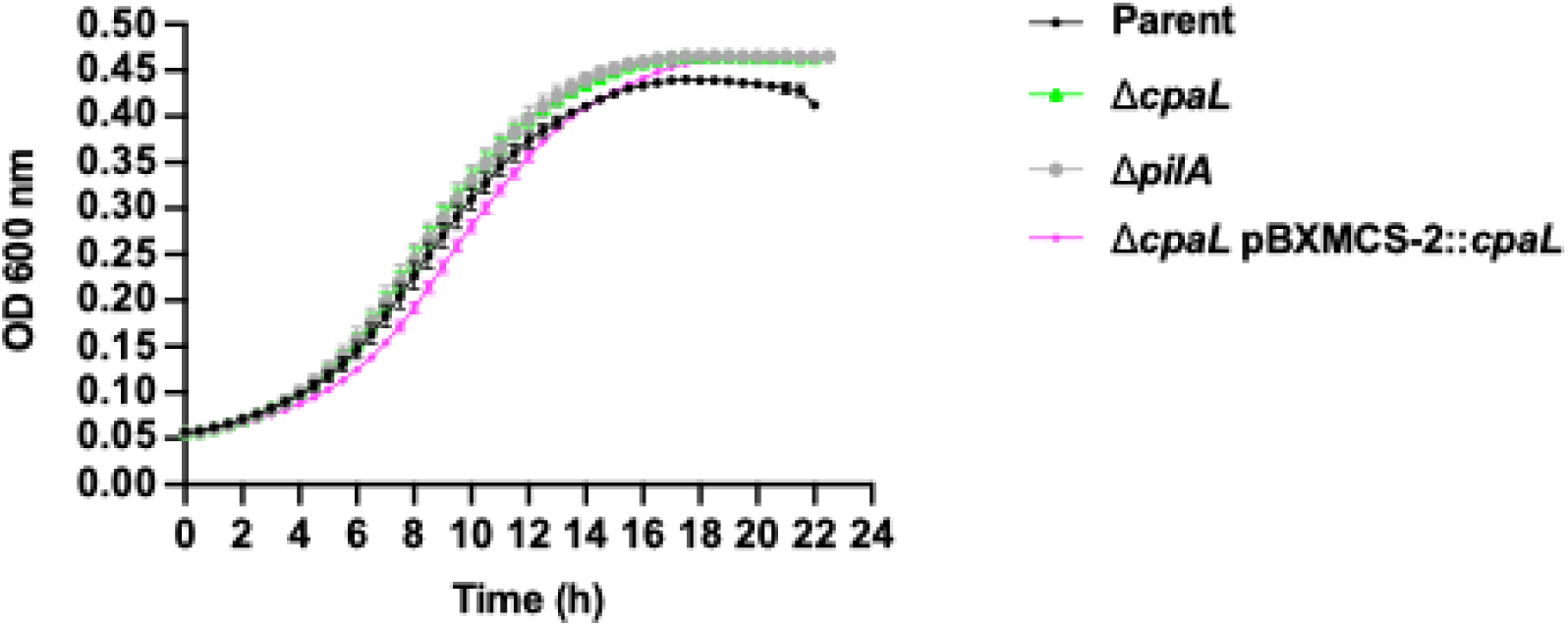
*Caulobacter* cell growth is not affected by the absence of *cpaL*. Growth curves of the indicated strains of *C. crescentus* grown in PYE. Data were collected every 30 min and are the average of six independent biological replicates. Error bars represent the standard error of the mean.

**Figure S3:**
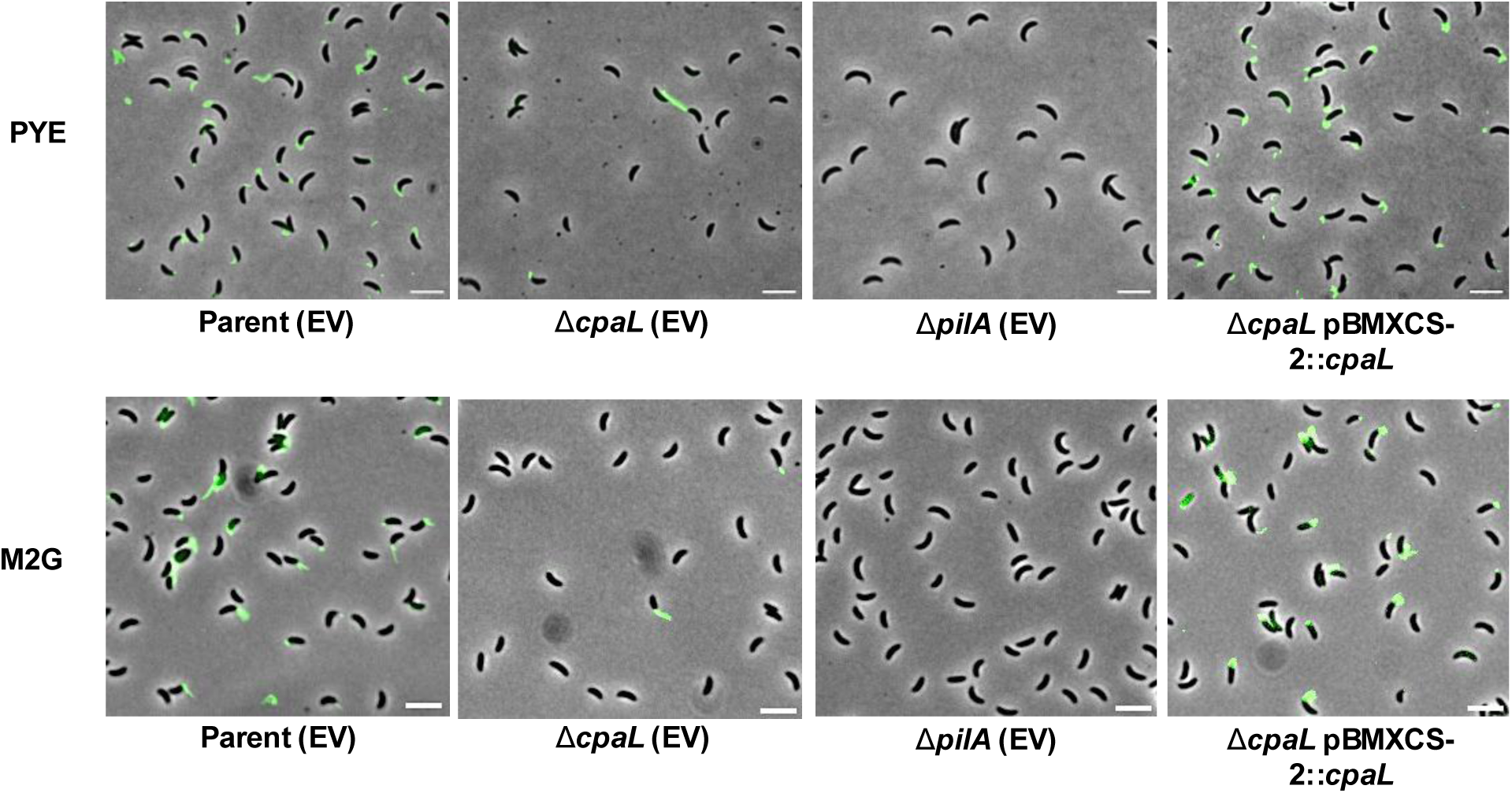
Representative microscopy images for the strains used in the phage assay in PYE and M2G medium. Synchronized swarmer cells were blocked for pilus retraction with PEG5000-mal and labeled with AF488-maleimide (green). Tested strains included the parent (EV: empty vector), the Δ*cpaL* (EV) mutant, the Δ*pilA* (EV) mutant, and the plasmid-complemented Δ*cpaL* mutant. Scale Bar, 10 μm.

**Figure S4:**
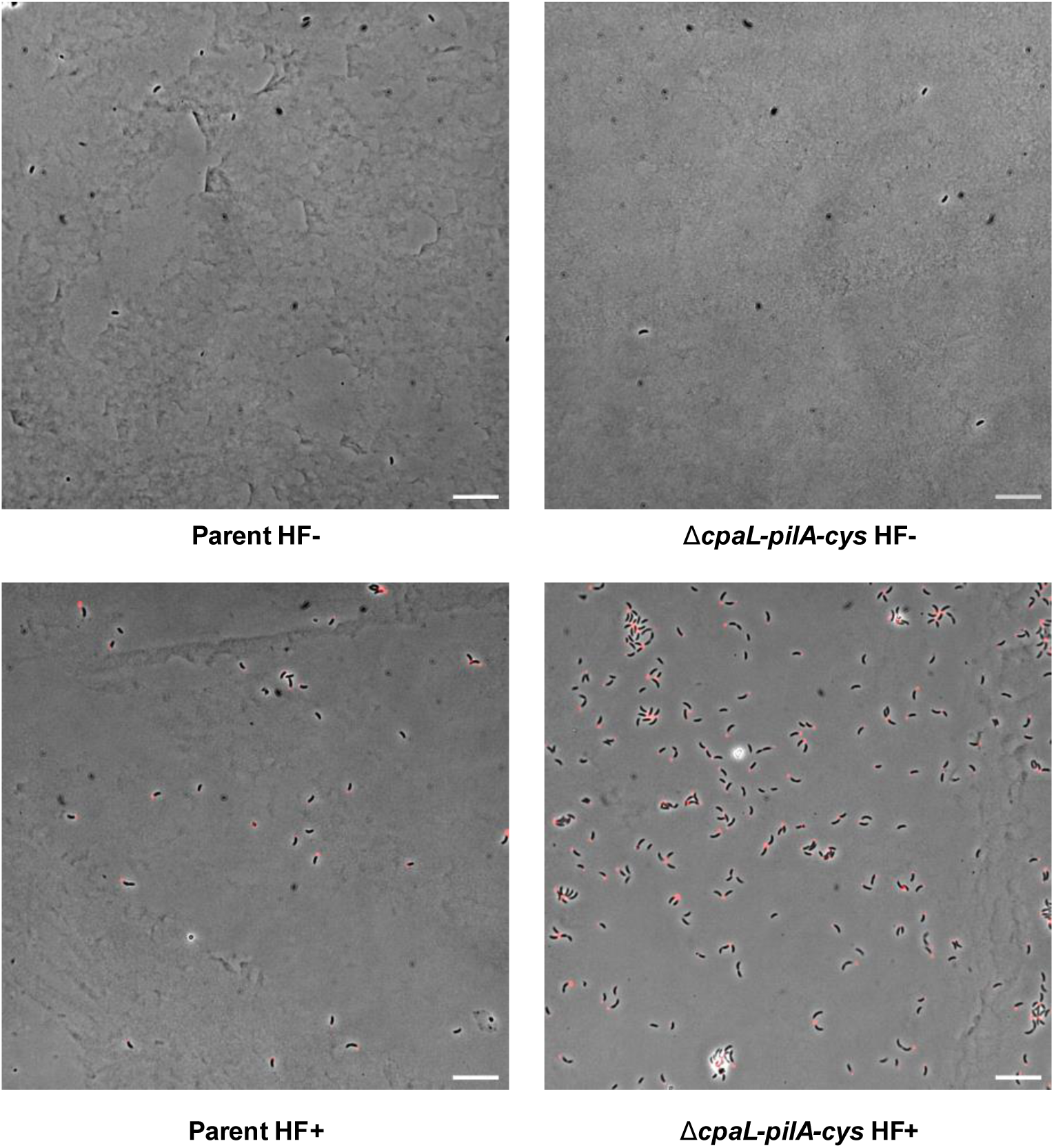
Representative microscopy images for cells attached to the surface. Cells are grown in the complex medium PYE and incubated for 30 min to a glass coverslip (HF+: Holdfast positive). Holdfasts are labeled with the AF488 conjugated wheat germ agglutinin (AF488-WGA). Scale Bar, 10 μm.

**Figure S5:**
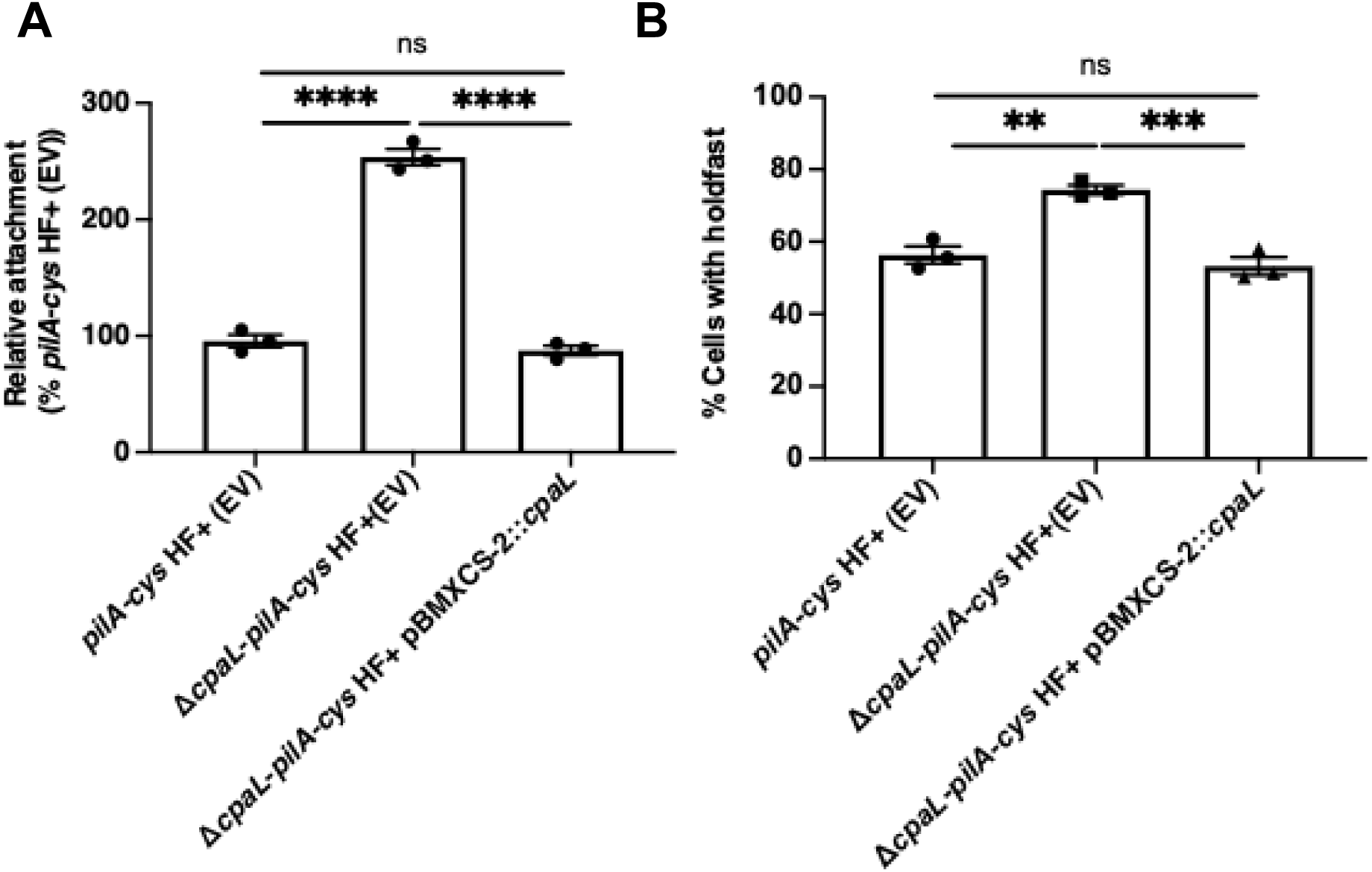
Plasmid complementation of the Δ*cpaL* mutant rescues its increased attachment and holdfast production phenotypes. **A:** Quantification of the attachment of cells grown in the complex medium PYE to a glass coverslip after 30 min of incubation **B:** Quantification of the percentage of cells producing holdfast in the population in the complex medium PYE on an agarose pad. Data are the mean of three independent biological replicates. Error bars indicate the standard error of the mean. Statistical comparisons were made using Tukey’s multiple comparisons test. ^****,^ P <0.0001; ^***,^ P <0.001; ^**,^ P <0.01.

**Figure S6:**
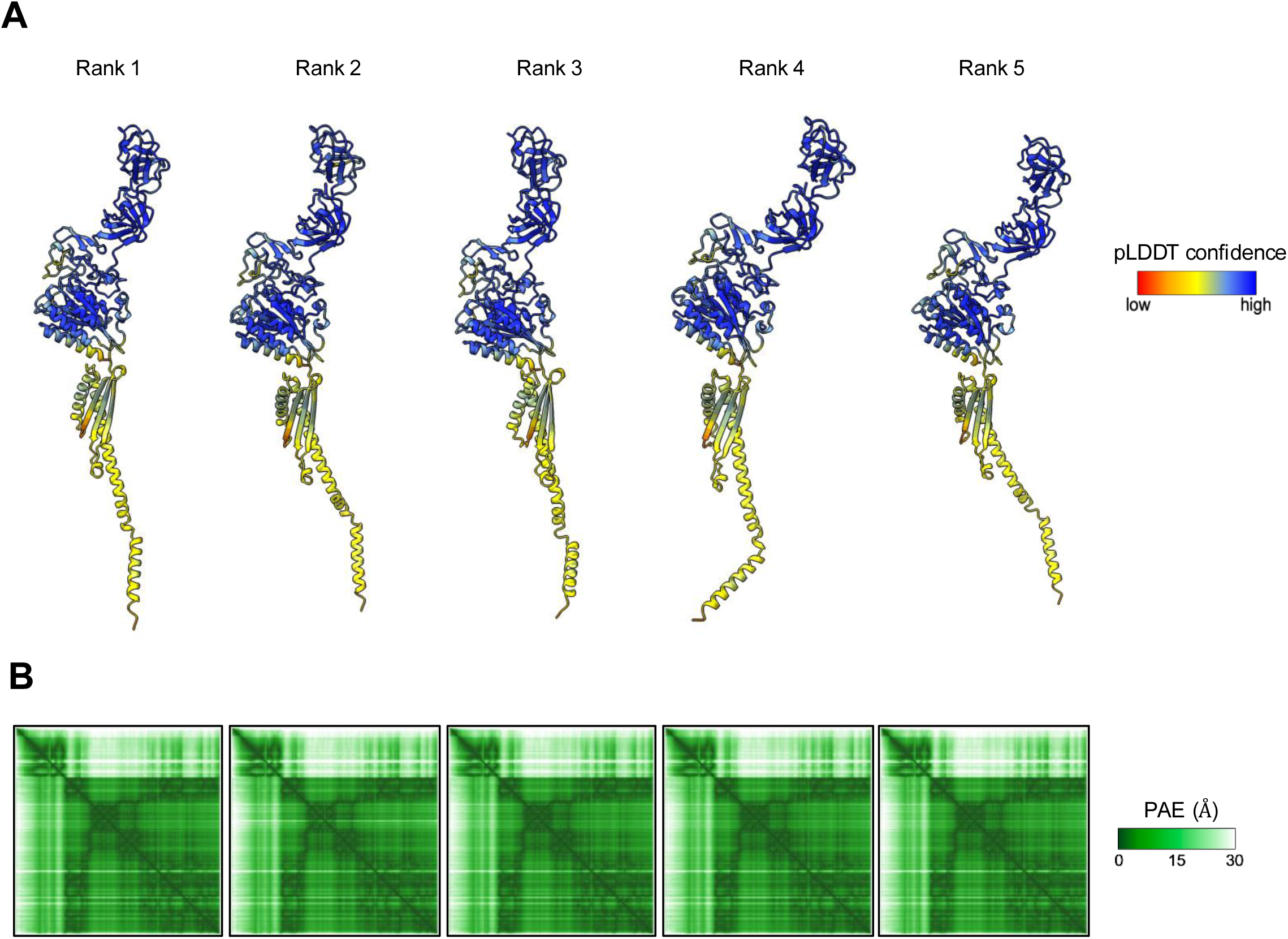
**A:** Representative of the five predicted model structures of CpaL predicted by Alphafold3. Structures are ranked according to the predicted template modeling (pTM) score and are colored according to the predicted local distance difference test (pLDDT) score, which indicates per-residue model confidence for the protein chain. The colors represented the measurement of the confidentiality in the prediction of the structure and are according to the predicted local distance difference test (pLDDT) score. **B:** Confidence in the prediction of the structure is indicated by the predicted aligned error (PAE) scores, which indicate positional error in angstroms for a given pair of residues within the protein chain.

**Figure S7:**
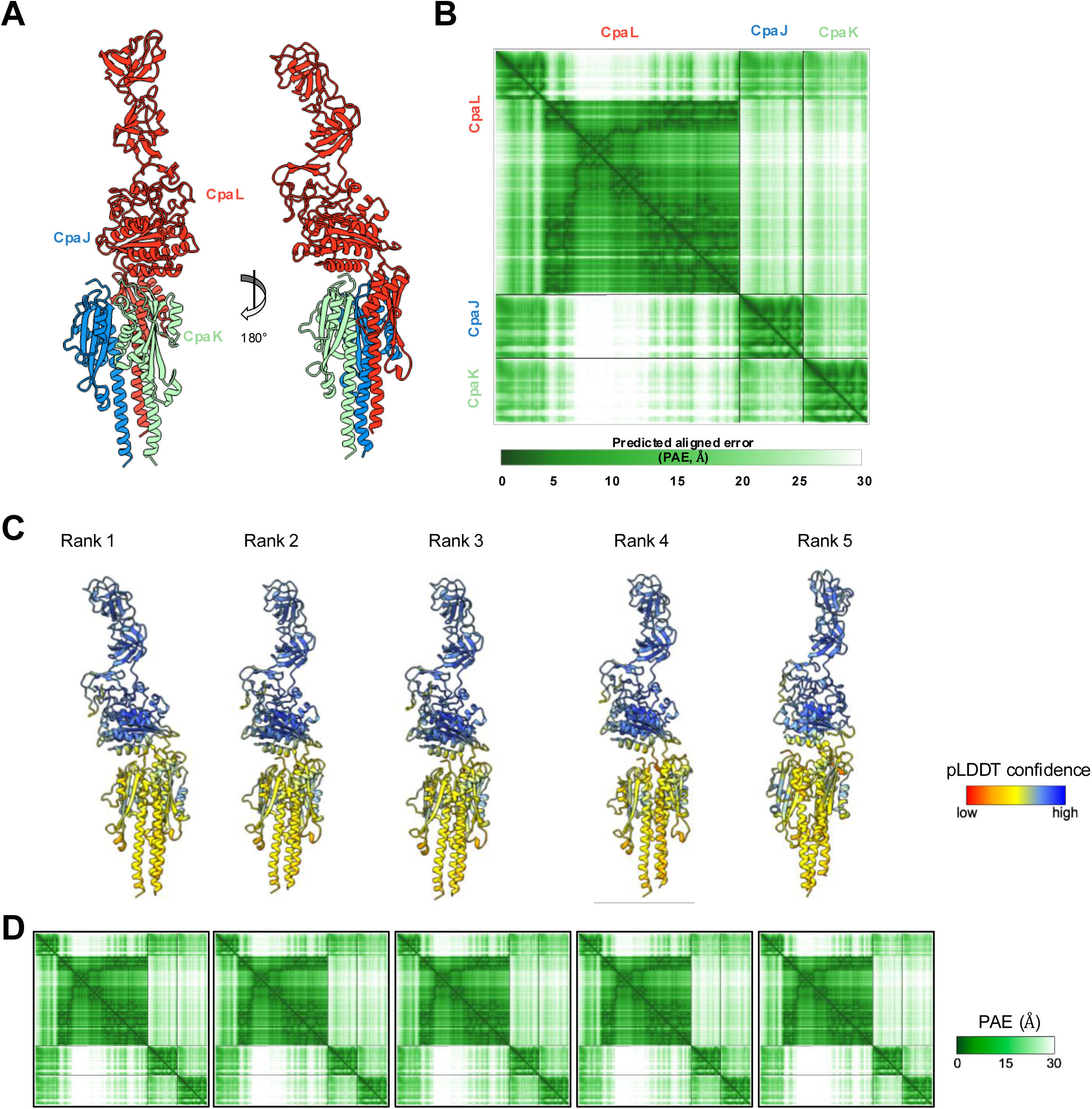
**A:** Top-ranked structure of a complex composed of the minor pilins CpaJ (blue), CpaK (green), and CpaL (red), predicted by AlphaFold3. The predicted or known signal sequences of each protein were removed prior to the prediction. **B:** The predicted aligned error (PAE) scores for the model depicted in panel A. The PAE indicates the positional error in angstroms (Å) for a given pair of residues across all protein chains in the model. **C:** Structures of a complex composed of CpaJ, CpaK, and CpaL, with their predicted signal sequences removed, predicted by AlphaFold3. Structures are aligned and shown from the same orientation. Structures are ranked according to the predicted template modeling (pTM) score and are colored according to the predicted local distance difference test (pLDDT) score, which indicates per-residue model confidence for the individual protein chains within the complex. **D:** Confidence in the prediction of the complex is indicated by the predicted aligned error (PAE) scores, which indicate positional error in angstroms for a given pair of residues across all protein chains.

**Figure S8:**
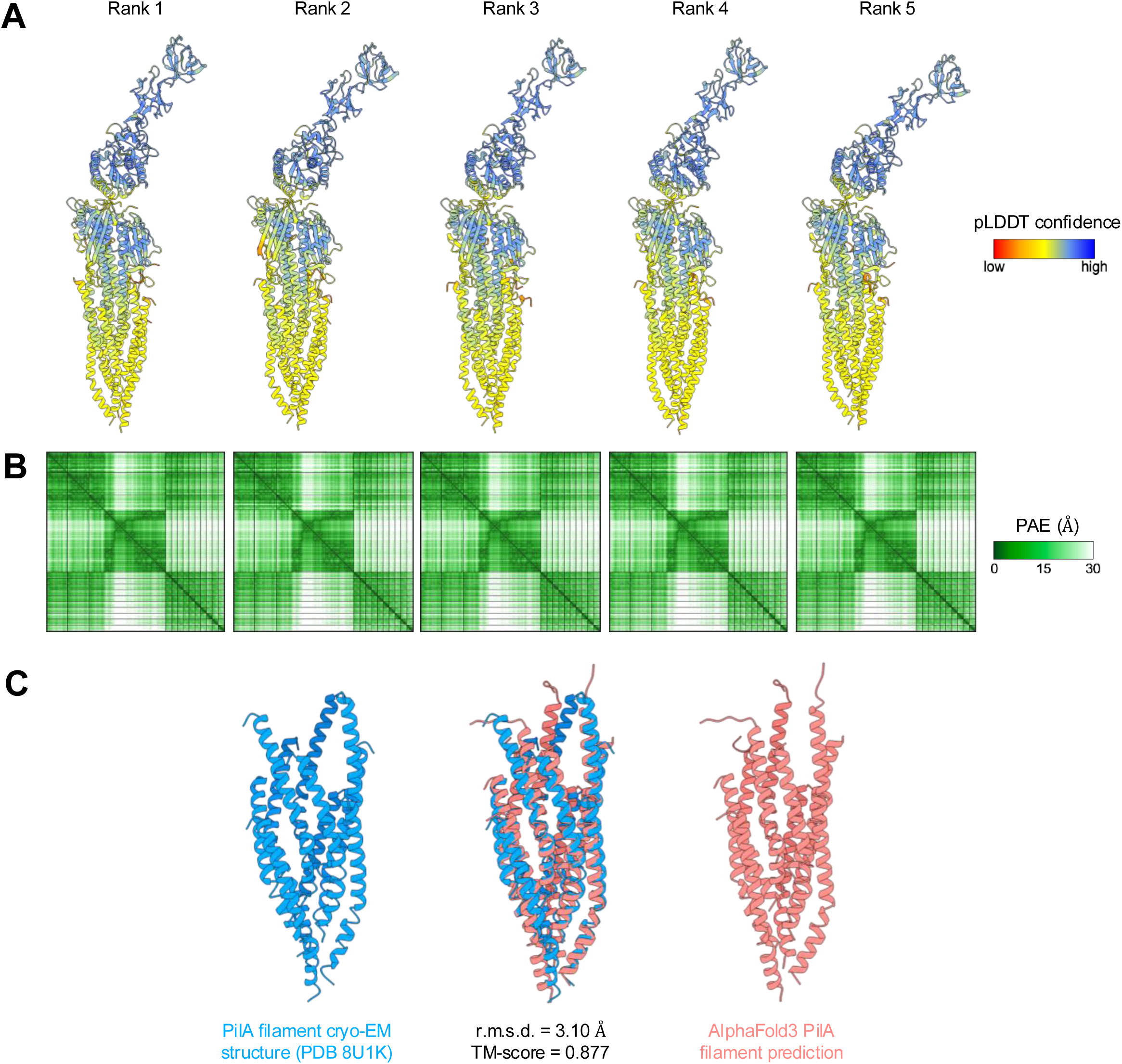
**A:** Structures of a complex composed of CpaJ, CpaK, CpaL, and ten PilA subunits, with their predicted signal sequences removed, predicted by AlphaFold3. Structures are aligned and shown from the same orientation. Structures are ranked according to the predicted template modeling (pTM) score and are colored according to the predicted local distance difference test (pLDDT) score, which indicates per-residue model confidence for the individual protein chains within the complex. **B:** Confidence in the prediction of the complex is indicated by the predicted aligned error (PAE) scores, which indicate positional error in angstroms for a given pair of residues across all protein chains. **C:** Alignment of the structure of the *C. crescentus* PilA filament determined by cryo-EM (from PDB 8U1K) and the structure of the PilA filament as determined by AlphaFold3 (from the prediction in panel A).

**Figure S9:**
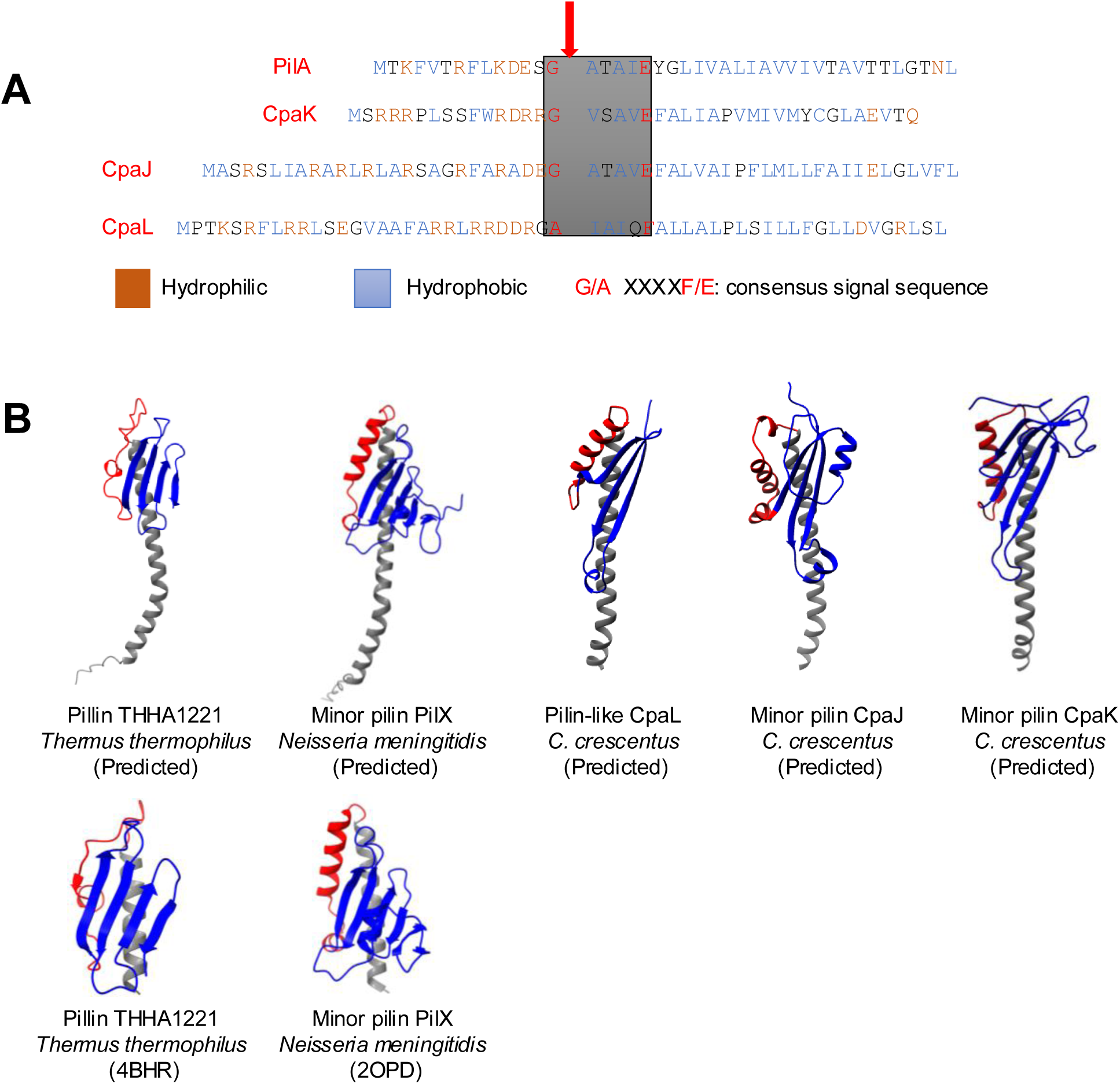
CpaL has a pilin-like module with a predicted prepilin peptidase (CpaA) cleavage site. **A:** N-terminal sequence of PilA, minor pilins CpaK and CpaJ, and CpaL. Hydrophilic residues are highlighted in orange, and hydrophobic residues are highlighted in blue. The consensus sequence (G/A-X-X-X-F/E) for recognition by the prepilin peptidase CpaA is shown in a grey box. The potential CpaA cleavage site is indicated with a red arrow. **B:** Structural comparison of the predicted pilin-like module of CpaL, CpaJ, and CpaK, with their predicted signal sequences removed, and the predicted and experimentally determined structure of the T4P THHA1221 from Thermus thermophilus (PDB:4BHR) and the minor pilin PilX from *Neisseria meningitidis* T4aP (PDB ID: 1AY2).

**Figure S10:**
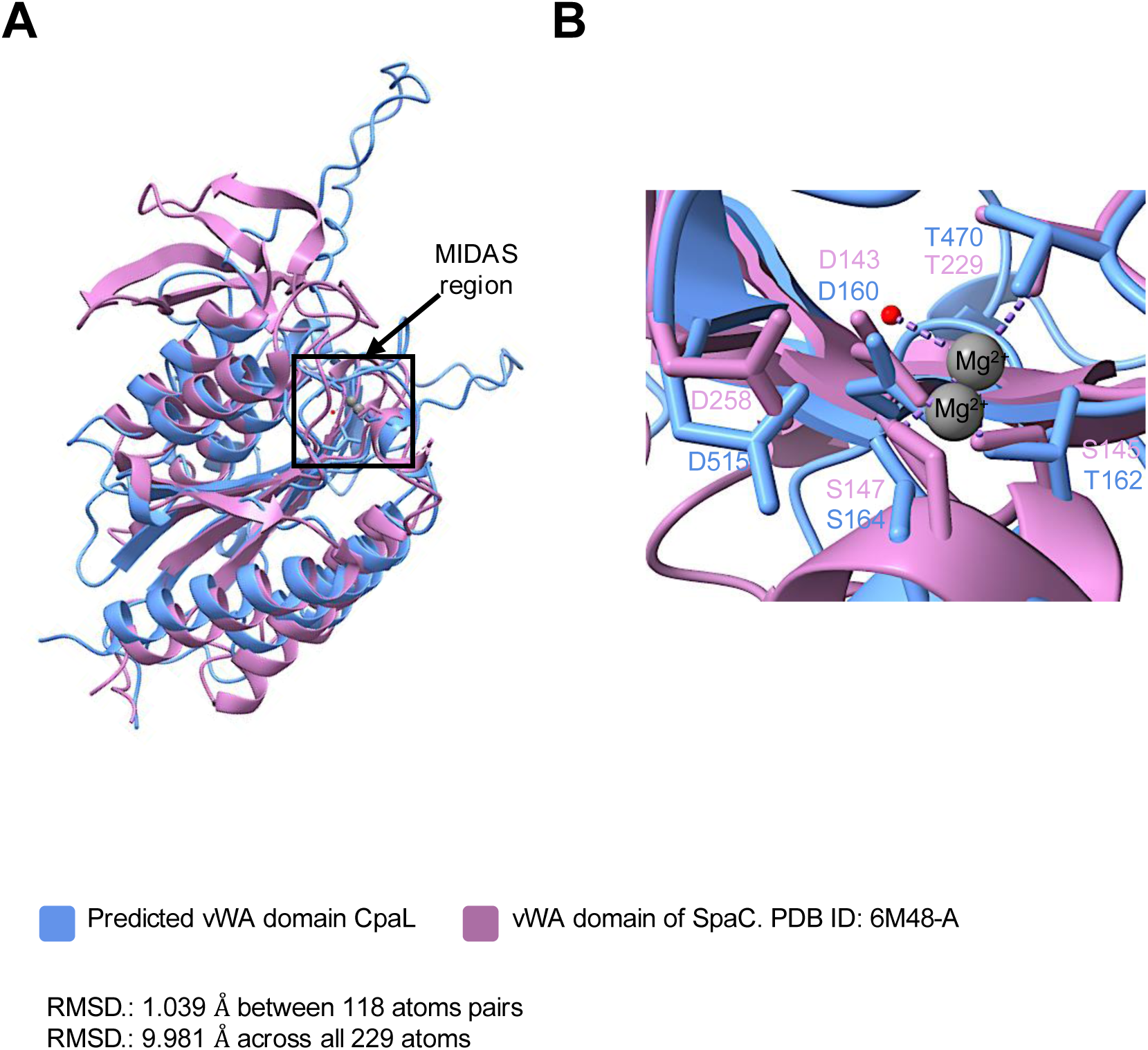
Structural comparison of the vWA domain of CpaL and the SpaC protein of *Lactobacillus rhamnosus* GG. **A:** Structural superposition of the predicted vWA domain of CpaL (blue, residues 148-203, and 383-626) with the vWA domain of SpaC (pink, residues 129-386) determined by X-ray crystallography (PDB ID: 6M48-A). **B:** Magnified image of the box in panel A depicting the MIDAS motif regions of CpaL (blue) and SpaC (pink). The Mg^2+^ ion that co-crystallized with SpaC as well as the Mg^2+^ ion predicted by Alhphafold3 for the vWA domain of CpaL are represented by two large grey spheres, while the water molecule involved in Mg^2+^ ion coordination by the SpaC are depicted as a small red sphere. RMSD: 1.039 Å between 118 atom pairs; RMSD.: 9.981 Å across all 229 atoms.

**Table S1:** Strains used in this study.

| Strain designation | Genotype and method of construction | Source or reference |
| --- | --- | --- |
| *Caulobacter crescentus* |  |  |
| YB8220 | NA1000 *hfsA*+ *pilA*<sup>T36C</sup> | (11) |
| YB6374 | NA1000 *hfsA*+ Δ*pilA* | (11) |
| YB8644 | NA1000 *hfsA*+ *pilA*<sup>T36C</sup> Δ*cpaL*<br>(Transduced strain YB8215 with lysate<br>made from strain YB8287) | This study |
| YB8454 | NA1000 *pilA*<sup>T36C</sup> Δ*cpaK* (Electroporated<br>plasmid pNPTS138Δ*cpaK* from strain<br>8453 into strain 8288. | This study |
| YB10278 | NA1000 *pilA*<sup>T36C</sup> Δ*cpaJ* (Electroporated<br>plasmid pNPTS138Δ*cpaJ* from strain<br>10277 into strain 8288. | |
| FC764 | NA1000 *hfsA*+ | (58) |
| YB8215 | NA1000 *hfsA*+ Δ*cpaL* (electroporated<br>plasmid from strain YB8206 into strain<br>FC764) | This study |
| YB8288 | NA1000 *pilA*<sup>T36C</sup> | (11) |
| LS3118 | NA1000 Δ*pilA* | (59) |
| YB8294 | NA1000 *pilA*<sup>T36C</sup> Δ*cpaL* (Electroporated | This study |
|  | strain YB8288 with plasmid from strain Y8206) |  |
| YB9770 | NA1000 *pilA*<sup>T36C</sup> pBXMCS-2<br>(Electroporated strain YB8288 with plasmid pBXMCS-2) | This study |
| YB9771 | NA1000 Δ*pilA* pBXMCS-2<br>(Electroporated strain LS3118 with plasmid pBXMCS-2) | This study |
| YB9772 | NA1000 *pilA*<sup>T36C</sup> Δ*cpaL* pBXMCS-2<br>(Electroporated strain YB8294 with plasmid pBXMCS-2) | This study |
| YB9742 | NA1000 *pilA*<sup>T36C</sup> Δ*cpaL* pBXMCS-2<br>*cpaL* (YB9741 Electroporated strain YB8294 with plasmid from YB9741) | This study |
| YB9773 | NA1000 *hfsA*<sup>+</sup> *pilA*<sup>T36C</sup> pBXMCS-2<br>(Electroporated strain YB8220 with plasmid pBXMCS-2) | This study |
| YB9774 | NA1000 *hfsA*<sup>+</sup> *pilA*<sup>T36C</sup> Δ*cpaL* pBXMCS-2<br>(Electroporated strain YB8644 with plasmid pBXMCS-2) | This study |
| YB9775 | NA1000 *hfsA*<sup>+</sup> Δ*pilA* pBXMCS-2<br>(Electroporated strain YB6374 with plasmid pBXMCS-2) | This study |
| YB9776 | NA1000 *hfsA*+ *pilA*<sup>T36C</sup> Δ*cpaL* pBXMCS-2 *cpaL* (Plasmid from YB9741 electroporated into strain YB8644 by electroporation) | This study |
| YB9777 | NA1000 *hfsA*+ *pilA*<sup>T36C</sup> Δ*cpaJ* (Electroporated strain YB 8220 with plasmid pNPTS138 Δ*cpaJ* from YB10277) | This study |
| YB9778 | NA1000 *hfsA*+ *pilA*<sup>T36C</sup> Δ*cpaK* (Electroporated strain YB 8220 with plasmid pNPTS138 Δ*cpaK* from YB 8453) | This study |
| *Escherichia coli* |  |  |
| YB4030 | S17-1/pNPTS138 Δ*pilA* | (19) |
| YB8206 | α-select/pNPTS139 Δ*cpaL* | This study |
| YB8286 | α-Select/pNPTS139 *pilA*<sup>T36C</sup> | (11) |
| YB8453 | α-select/pNPTS138 Δ*cpaK* | This study |
| YB102777 | α-select/pNPTS138 Δ*cpaJ* | This study |
| YB9741 | NEB 5-alpha/pBXMCS-2 *cpaL* | This study |

**Table S2:**
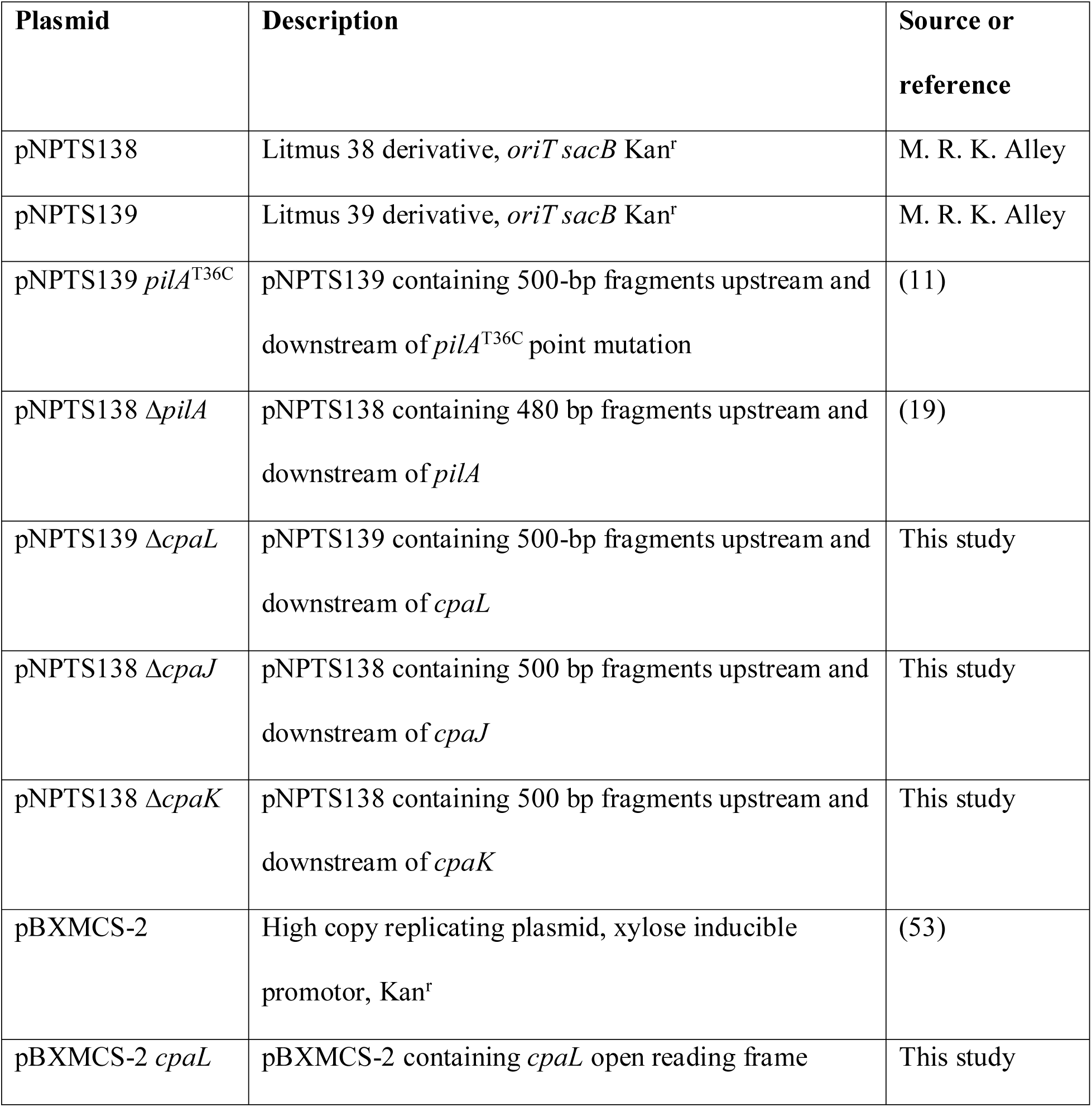
Plasmids used in this study.

**Table S3:**
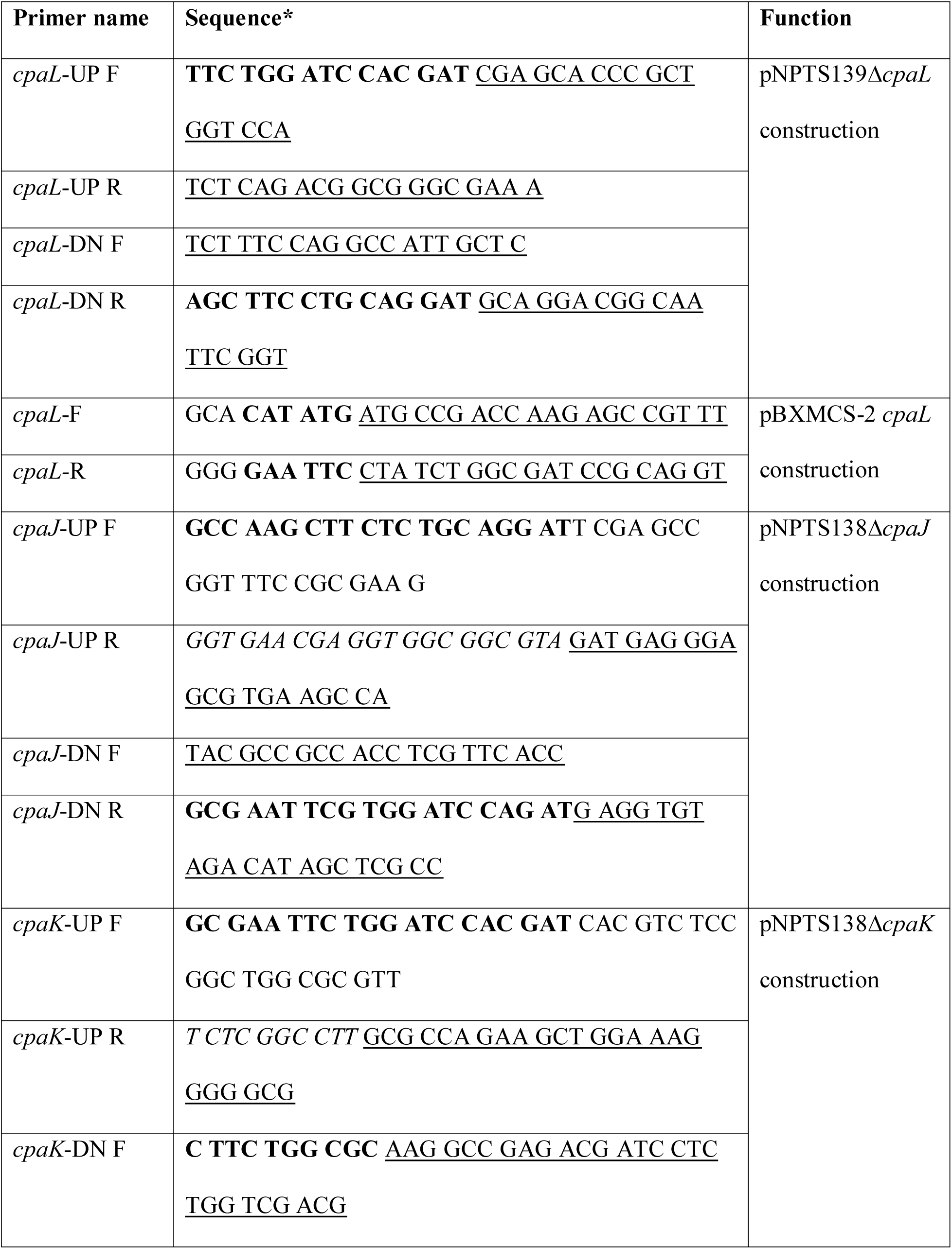
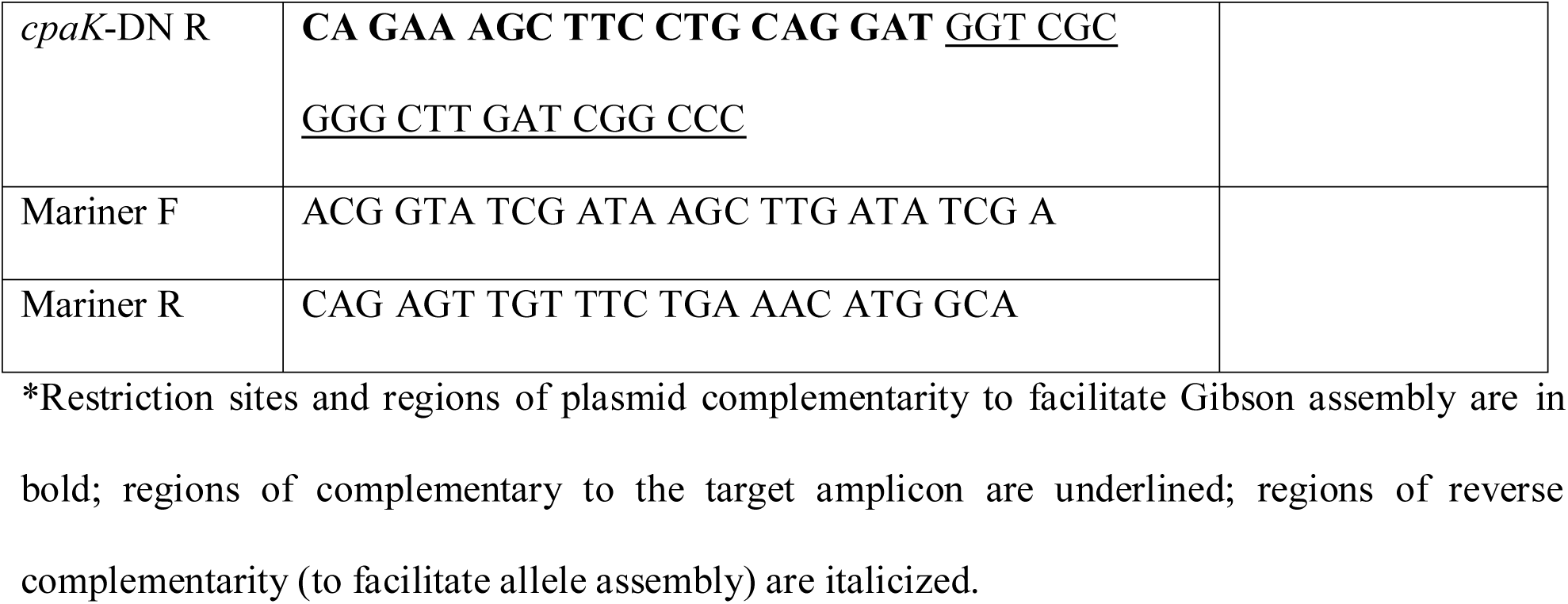
Primers used in this study.

**Table S4:** The top five PDB hits identified by the DALI server for the predicted structure of CpaL.

| PDB hit | Description | Species | Z-score | rmsd | lali | nres | %id |
| --- | --- | --- | --- | --- | --- | --- | --- |
| 6m48-A | SpaC | *Lactobacillus rhamnosus GG* | 20.3 | 4.1 | 241 | 813 | 16 |
| 2ww8-A | Cell wall surface anchor family protein; Pilus adhesion RrgA | *Streptococcus pneumoniae* | 19.9 | 13.2 | 254 | 815 | 18 |
| 6to1-A | Minor fimbrium subunit Mfa5 | *Porphyromonas gingivalis* | 18.9 | 4 | 232 | 566 | 21 |
| 7b7p-A | Type IV pilus biogenesis protein PilB | *Streptococcus sanguinis* | 18.7 | 4.8 | 244 | 420 | 17 |
| 7w6b-A | von Willebrand factor type A domain protein | *Streptococcus oralis* | 18.3 | 5.5 | 212 | 782 | 21 |

Rmsd: Root mean square deviation, calculated across only the aligned portions of CpaL and the hit; lali: The number of residues of the hit protein that were aligned with CpaL; nres: the total number of residues in the hit protein; %id: percentage identity between CpaL and the hit protein.

## Notes

### Competing Interest Statement

The authors have declared no competing interest.

### Summary of Updates

Supplementary movies uploaded. Four supplementary movies have been uploaded to this manuscript.

